# Tension mediated mechanical activation and pocket delipidation lead to an analogous MscL state

**DOI:** 10.1101/2021.04.01.438050

**Authors:** Bolin Wang, Benjamin J. Lane, Charalampos Kapsalis, James R. Ault, Frank Sobott, Hassane El Mkami, Antonio N. Calabrese, Antreas C. Kalli, Christos Pliotas

## Abstract

The MscL channel gates in response to membrane tension changes to allow the exchange of molecules through its pore. Lipid removal from transmembrane pockets leads to a MscL response. However, it is unknown whether there is correlation between the tension mediated state and the state derived by pocket delipidation in the absence of tension. Transitions between MscL states may follow a similar pathway to cover the available conformational space but may not necessarily sample the same discrete intermediates. Here, we combined pulsed-EPR and HDX-MS measurements on MscL, coupled with molecular dynamics under membrane tension, to investigate the changes associated with the distinctively derived states. Whether it is tension or pocket delipidation, we find that MscL samples a similar expanded state, which is the final step of the delipidation pathway but only an intermediate stop of the tension mediated path. Our findings hint at synergistic modes of regulation in mechanosensitive channels.

## Main Text

### Introduction

Specific lipid binding events and changes in transbilayer pressure can modulate the structure of membrane proteins and regulate their function (1–11). Mechanosensitive (MS) ion channels are membrane proteins which sense and respond to tension changes in the cell membrane. Pressure-sensitive domains, or pockets, formed within transmembrane (TM) regions of bacterial MS channels have been proposed to play a role in their mechanical sensing and response (1, 2, 8, 12, 13). These pockets were first identified on the MS channel of small conductance MscS, leading to the lipids-move-first hypothesis, which predicts that lipids accessing the pockets physically inhibit (directly or allosterically) the movement of pore helices to gate the channel (2). When a modification (L89W) was introduced at the entrance of these pockets in MscL, steric lipid interactions with the backside of the pore-lining helix TM1 were significantly reduced and as a consequence MscL expanded, adopting an intermediate state, even in the absence of applied tension(8). MscL’s mechanical response suggested that pocket targeting lipids could act as negative allosteric modulators for MS channels and that tension could be mimicked by molecules or gain-of-function modifications targeting the pocket region, which could disrupt the lipid pathway between the bulk membrane and the pockets to gate the channel (1, 8, 14, 15). Indeed, the function of MscL and MscS (5, 14) can be modulated by changes in molecule occupancy within these pockets (2, 8, 16–18) and even subtle structural differences within the same regions could lead to functional differences in different MscL proteins(19). This suggests a common regulatory mechanism, tailored to the unique structural landscape of MS channels, which may guide ligand specificity.

In mammals, the MS channel Piezo1 is activated by Yoda1, opening the possibility of endogenous agonists(20, 21), while TRAAK and TREK-1 are responsive to natural chemical ligands including arachidonic acid, polyunsaturated fatty acids, and lysophosphatidic acid(22–24). Lipids residing within similar inner-leaflet pockets have also been linked with mechanosensation in x-ray and cryo-EM structures of TRPV3(25), TRAAK(26), TREK-2(27), YnaI(28), MSL1(29) and MscS(2, 17, 30, 31), while their unique parallel to membrane plane orientation may be guided by an amphipathic helix that is important in the mechanosensitivity of MscL(32) and GPCRs(33). Pulsed electron-electron double resonance (PELDOR, also known as DEER)(34, 35) and x-ray studies on the LmrP multidrug transporter showed that lipids could provide a hydrophobic counterpart to partially fill binding pockets and participate in the coordination of ligand binding(36).

Attempts to model MscL opening have been previously implemented via steered MD simulations, with added tensions to pull the channel open (37–41), but the tension used in this modelling is an order of magnitude higher than that required to open MscL in giant unilamellar vesicles (GUVs) and spheroplasts(8, 42, 43). In other cases, MscL was subjected to extensive modifications or forces were applied for relatively short periods of time(37, 44). MD simulations on TREK-2 have shown that under increased bilayer tension the channel moves from a “down” to an “up” configuration accompanied by cross sectional area changes, while its pockets are occupied by lipids only in the absence of membrane stretch(45).

Hydrogen Deuterium eXchange Mass Spectrometry (HDX-MS) and 3 pulse Electron Spin Envelope Echo Modulation (3pESEEM) spectroscopy are powerful tools in the investigation of membrane protein structural dynamics. The former involves the exchange of protons on a protein with deuterium to generate accessibility information, and has been used to provide key insights into the role(s) of lipids in modulating membrane protein conformation(46–49). The latter measures weak hyperfine coupling between unpaired electrons and nuclear spins to probe accessibility to solvent (D_2_O) and has been used to investigate the dynamics of membrane proteins(8, 19, 34, 35, 50–54). Here we endeavored to investigate whether there is a structural analogy between the physiologically relevant tension-activated state, and the one stabilized by modifications that result in pocket lipid removal. To this end, we have combined untargeted (HDX-MS) and single-residue (3pESEEM) methods to probe the architecture of MscL upon pocket delipidation and independently generate a tension-activated MscL state by MD simulations. We find these two differently derived states lead to an analogous state, suggesting a direct link between tension- and pocket lipid removal- activation in MS channels. Further, our data suggest that upon tension-activation additional structural changes can trigger substantial further opening of the channel pore, suggesting that structural plasticity of these mechanosensitive ion channels enables them to respond differently upon receipt of discrete stimuli.

### Results

#### Channel gating by lipid pocket removal activation monitored by HDX-MS

The substitution of a tryptophan at position L89 in *Mycobacterium tuberculosis* MscL (TbMscL) has been shown to restrict lipid access to channel pockets and destabilise the closed state, leading to an expanded (sub-conducting) MscL state(8). In order to give a mechanistic insight into the transitions occurring between the two states, we used HDX-MS to measure relative differences in deuterium uptake between the WT (closed) and L89W (expanded by pocket delipidation) channel proteins. We succeeded in obtaining 95% peptide coverage of the entire resolved MscL structure where the gating occurs, i.e., residues 1-125, including the entire TM domain and the largest part of the cytoplasmic helical bundle (83% overall coverage for the entire construct including the C-terminal 6xHis-tag). Differences in uptake, detected at the peptide level, between the two states allowed us to identify protein regions that became deprotected from exchange in the sub-conducting L89W state (Fig 1). Three regions of MscL were identified and showed significant changes between the two states. Peptides in the regions containing residues 37-53 (periplasmic loop), 58-69 (top of TM2), and 97-111 (bottom of TM2, cytoplasmic loop and top of the cytoplasmic helical bundle) had a significantly higher uptake of deuterium in L89W in comparison to WT TbMscL (Fig 1, S1 and Table S1). Residues 37-53 run from the top of TM1 and end after the first β-sheet in the loop connecting TM1 to TM2. Residues 58-69 correspond to the middle of the periplasmic loop up until TM2, and residues 97-111 cover the bottom of TM2 to the top of the C-terminal helical bundle (Fig 1 and S1). This gave us the first glimpses into MscL’s pocket lipid removal activation and, which protein domains are mostly impacted by the L89W (pocket entrance) modification across the entire MscL’s length.

**Figure 1.**
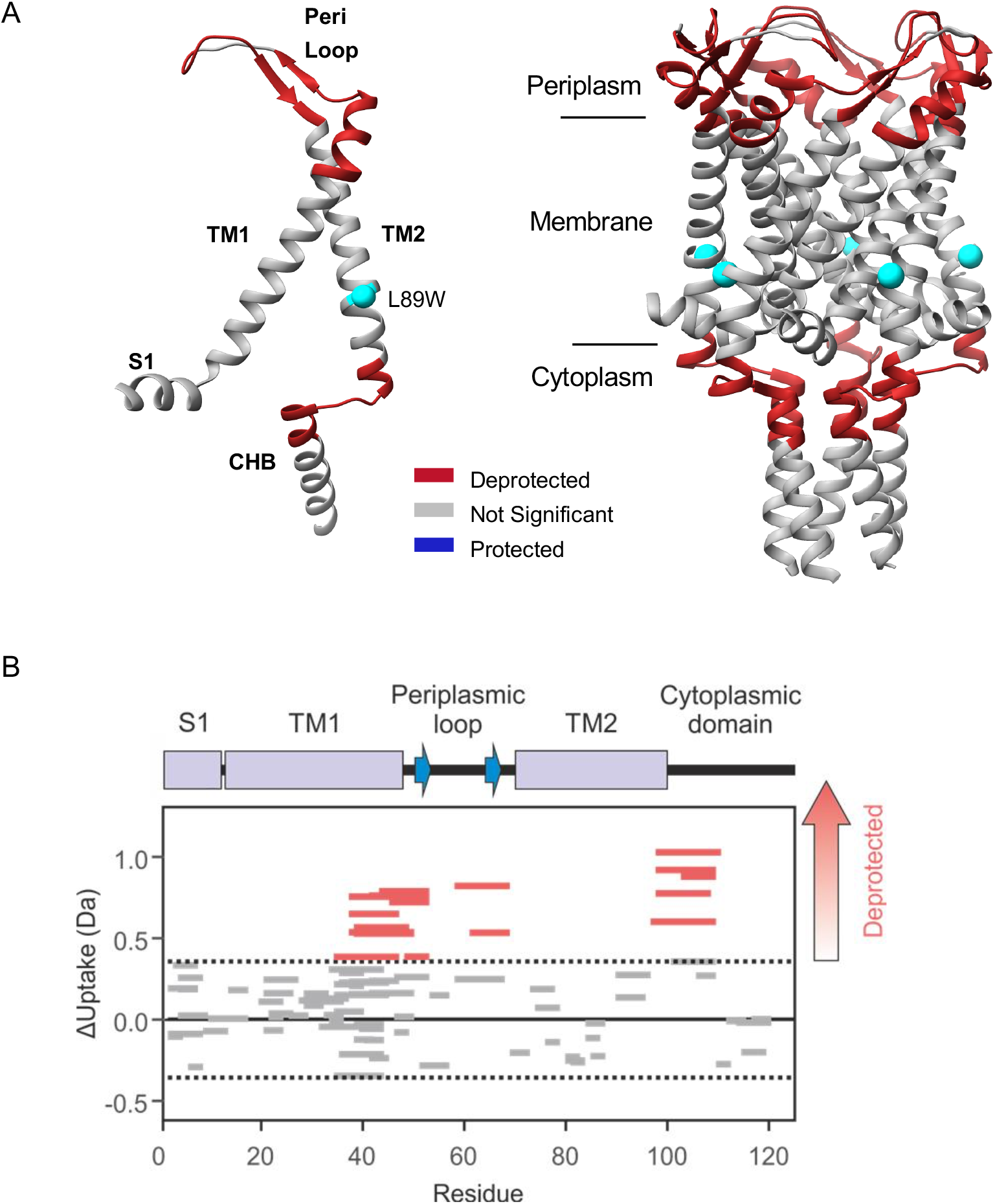
Global changes on MscL induced by pocket lipid removal observed by HDX-MS. A. Differences in the deuterium uptake of regions of TbMscL (PDB 2OAR) when comparing the WT and the L89W (pocket lipid removal) modified protein. Regions highlighted in red are deprotected following the L89W modification. Regions of the protein in grey show no significant difference between the two conditions. L89W modification site is depicted as a cyan sphere. B. Wood’s plots showing the summed differences in deuterium uptake in MscL over all five HDX timepoints, comparing WT with L89W MscL (Wood’s plots were generated using Deuteros(79). Peptides coloured in red, are deprotected from exchange in L89W MscL. No peptides were significantly protected from exchange in L89W MscL compared with wild type MscL. Peptides with no significant difference between conditions, determined using a 99% confidence interval (dotted line), are shown in grey. Example deuterium uptake curves are shown in Fig S1.

#### Solvent accessibility mapping at single residue resolution by 3pESEEM

Next, we sought to investigate the feasibility of 3pESEEM to study the structure of MscL and identify sites that are potentially dynamic during gating and solvent accessible. To this end, we generated twenty-six single cysteine mutants, accounting for 20% of the total TbMscL length, spanning all protein domains. Specifically, L2 and F5 on the S1 helix, N13, L17, V21, I23, V31 and F34 on TM1, L42, and F48 on the periplasmic loop, residues N70, V71, L72, L73, S74, F79, F84, A85, Y87, F88, L89, R98, K99, and K100 on TM2, and E102 and V112 on the cytoplasmic loop and helical bundle (CHB) respectively, were mutated to Cys in a Cys-free WT TbMscL background (Fig 2). We subsequently expressed, purified and spin labelled each one of these twenty-six mutants (MTSSL modification is denoted as R1 hereafter) and performed 3pESEEM solvent accessibility measurements (Fig S2). Solvent accessibility is based on the modulation depth of deuterium in the time domain signal, which is proportional to its associated signal intensity in the frequency domain. We used two independent analysis methods to determine solvent accessibility, both yielding very similar results (Fig S3). Unlike PELDOR which requires high spin labelling efficiency to obtain high sensitivity data acquisition(8, 13), 3pESEEM can be performed with no major losses in sensitivity even if the sample of interest shows lower spin labelling efficiencies for individual sites. This allowed us to obtain good quality spectra and quantify the solvent (D_2_O) accessibility for all twenty-six sites (Fig 2, S2 and S4). Residues in the S1 helix showed low accessibility, which is consistent with previous reports suggesting that this helix is buried within the bilayer(32). TM1 pore-lining residues presented low to intermediate accessibility, as would be expected for a closed channel pore(55). I23, V31 and F34 on TM1 are intermediately solvent-exposed (or relatively buried) and positioned on the same side of TM1 (Fig 2 and S4). This could be due either to the presence of native lipids, detergent used for membrane protein extraction and/or hinderance by the presence of TM2 on the TM1 interface. L42 and V48 are intermediately to highly exposed, as they form part of the periplasmic loop connecting TM1 and TM2. N70, V71, L72, L73, and S74 present the largest disparity across our whole data set. L72 is the most buried residue we measured (~ 3% compared to the most solvent-exposed cytoplasmic-facing labelled residue K100R1) while N70 (~90%) and V71 (~80%) are two of the most exposed (~90%), comparable to D_2_O (solvent) accessibilities measured for exposed cytoplasmic sites (Fig 2, S2 and S4). The remaining residues present intermediate accessibilities to these two extremes, as expected for subsequent residues, which form a helical turn with different space orientation. Residues N70 and V71 are solvent-exposed in the closed MscL state, while residues L72 and L73 are buried. The latter pair of residues is known to become exposed during opening, due to an anti-clockwise TM2 rotation(8, 19, 56). F79, F84 and A85 are relatively buried and lie at the interface of TM helices between different MscL subunits. Y87 has been identified as a lipid binding site(57) and along with F88 and L89 presented low to intermediate accessibilities and forms part of the pocket region(8, 19, 58). Except for R98, which showed intermediate accessibility, K99 and K100, presented high accessibilities, indicating these residues are solvent-exposed in the closed state. R98, K99 and K100 form a positively charged cluster for attracting negatively charged lipids(57) and our data suggest that the non-ionic DDM binds either differently or at a different site. Substitution of these residues with the neutral Gln had no effect on MscL’s conformation, suggesting that specific lipid headgroup binding on this particular site does not influence MscL gating(8). E102 (cytoplasmic loop) and V112, which is located in the upper portion of CHB and is expected to move apart upon MscL opening (59), presented high solvent accessibilities. Combined, this has enabled us to validate 3pESEEM as a suitable tool to map the structure of MscL.

**Figure 2.**
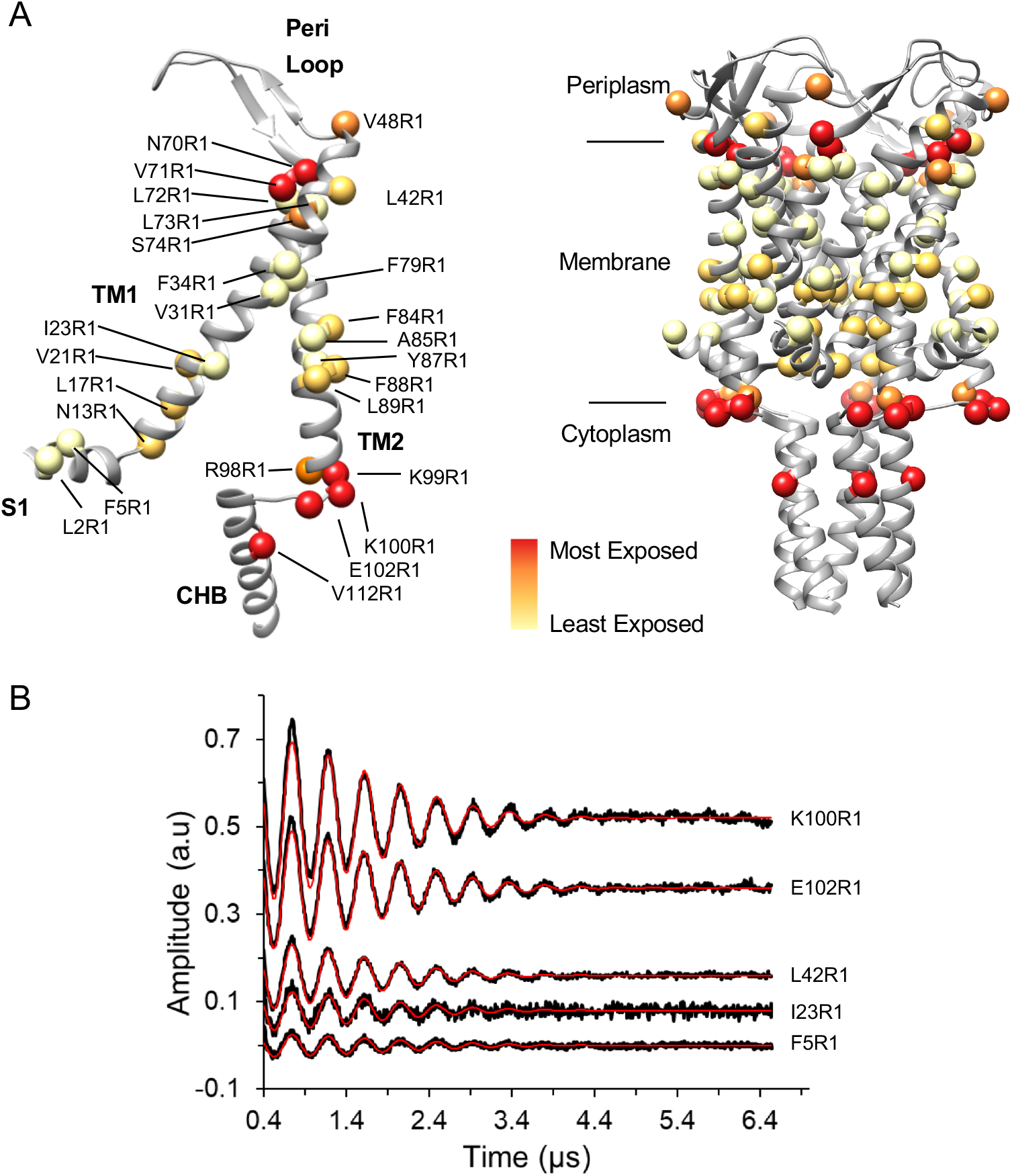
Single-residue mapping of MscL by 3pESEEM. A. Solvent accessibility of spin-labelled residues are shown as red and pale-yellow spheres for the most and the least solvent accessible, respectively. Spin-labelled residues are coloured based on the quartile of their relative accessibility which has been normalised on scale of 0% and 100%, where the residue with the highest accessibility corresponds to 100%. B. Background-corrected time-domain 3pESEEM experimental spectra with fitting for representative spin-labelled mutants. Residue F5R1 is found on the S1 amphipathic helix, I23R1 and L42R1 on TM1, and K100R1 and E102R1 are at the interface between TM2 and the CHB.

#### Identification of channel gating regions with single residue resolution

HDX-MS resolution is usually limited to the peptide level and, therefore, it is often unable to resolve changes that occur at different residues within the same peptide. Here we have already shown that the solvent accessibility of individual residues of MscL can be probed using 3pESEEM. This generates higher resolution information pertaining to the regions of structural change identified by HDX-MS, as well as to the channel pore vapour lock. In each of these regions, we introduced eight distinct single cysteine substitutions in an L89W MscL background for comparison with identical sites in the background of the wildtype protein. Specifically, N13 and V21 (pore), L42 in the periplasmic loop, N70, V71, L72, L73 (top of TM2) and K100 (bottom of TM2 and cytoplasmic loop) (Fig 3, S5 and S6). V21 forms the vapour lock (along with L17) that controls the channel gate and N13 sits one turn of a helix below. No significant change in this region was highlighted via HDX-MS, while due to the proximity of spin labels pointing towards the pore, distances are out of PELDOR’s distance measurement range (< 19 Å) (Fig 1 and 3B). However, 3pESEEM is highly sensitive to the local spin label environment, does not need a reference state as it is the case for HDX-MS, and is not subjected to distance restriction between sites like for PELDOR. Differences in the vapour-lock region may not have been detected by HDX-MS as structural changes may have resulted in complex changes to hydrogen bonding networks, whereby some residues were deprotected from exchange and others became protected, resulting in only minor/no changes in deuterium uptake at the peptide level (Fig 1 and S1). To resolve this, we labelled the V21 and N13 sites crucial for the channel’s vapour lock, and six other sites within the regions identified by HDX-MS, which undergo conformational changes when the L89W modification is present (Fig 3, S5 and S6). Overall, the motivation behind this was to confirm the peptide-level HDX-MS data and complement this with single residue resolution 3pESEEM data for crucial sites where changes may have not been detected, due to limitations associated with peptide-level analysis in HDX-MS.

**Figure 3.**
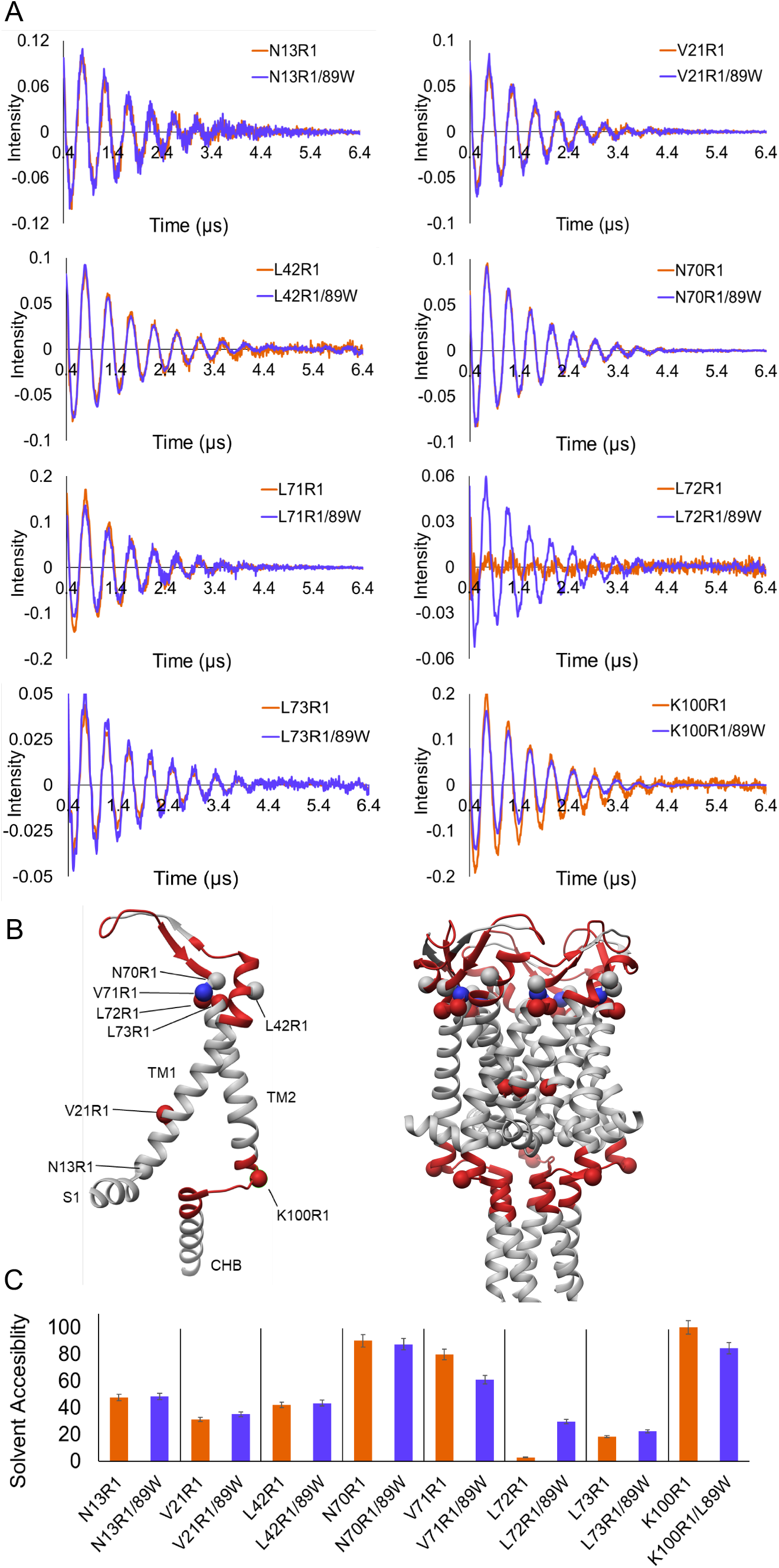
Effect of pocket delipidation on MscL structure investigated by integrated 3pESEEM and HDX-MS. A Background-corrected time-domain 3pESEEM experimental spectra (orange) of the single mutants N13R1, V21R1, L42R1, N70R1, L72R1, L73R1, and K100R1 in DDM overlaid with their associated L89W double mutant spectra (purple). B Differences in solvent accessibility for TbMscL (PDB 2OAR) following L89W modification. Spin labelled mutation sites used for 3pESEEM accessibility measurements are represented by spheres, and peptides that demonstrate a change in accessibility in HDX-MS following the L89W modification are represented as highlighted helices. Red regions or spheres highlight areas that are deprotected, while blue spheres and regions show areas that are protected following the L89W modification. C Column bar charts represent the 2H accessibility derived from the fitting of 3pESEEM time-domain traces. Errors are calculated conservatively at 5% to compensate for the fitting error and differences in relaxation times of different spin labelled residues. There was no significant difference in solvent accessibility of N13R1, L42R1, and N70R1 compared to their 89W double mutant counterpart. Solvent accessibility increased for V21R1, L72R1, and decreased for L71R1 and K100R1 following the L89W modification.

3pESEEM measurements at position L42, located in the periplasmic loop, showed no significant difference in solvent accessibility between the WT and the L89W states. For N13 which points towards the cytoplasm and is solvent-exposed in the closed state, we observed similar exposure in the expanded L89W MscL state (Fig 3 and S7). In L89W MscL, the solvent accessibility increased for V21, which is consistent with channel pore hydration as its side chain sits within the TM domain in the closed state (Fig S7). An over ten-fold increase in solvent accessibility was observed for L72, while a significant decrease was observed for K100, which sits at the C-terminal end of TM2 (Fig 3, S5 and S6). A decrease in solvent accessibility was observed for N70 and V71 in L89W MscL with 2.4% and ~19% reduction, respectively. Despite being consecutive residues, 70-73 show distinct changes in solvent accessibility in 3pESEEM, unlike HDX-MS, which reported an average difference in deuterium uptake at the peptide level. A significant and smaller increase in accessibility is seen for L72 and L73, which is opposite to the effect observed for N70 and V71, while the bottom of TM2 (K100) becomes less accessible to solvent in the L89W background (Fig 3, S5 and S6). This result is consistent with a rotation of the top of TM2 which occurs during mechanical activation and agrees with PELDOR distance measurements and x-ray structures(8, 19, 56). HDX-MS experiments showed deprotection on peptides inclusive of these residues and is consistent with a dramatic increase in accessibility of residue L72 upon pocket lipid removal activation (i.e. when the L89W mutation is present) in 3pESEEM measurements (Fig 3). V21 forms the vapour-lock for TbMscL and shows an increase in solvent accessibility in an L89W background, consistent with the pore gate hydration and the channel entering an expanded (and intermediate for full opening) state. HDX-MS measurements show a deprotection of the TM2 bottom for L89W versus WT suggesting an overall solvent exposure of this region (Fig 1 and 3). The MscL cytoplasmic loop has been shown to participate in gating and provide access to streptomycin entering the cell, while shortening of the loop influences MscL’s function by irreversibly decreasing its conductance(16, 60). Combined, HDX-MS and 3pESSEM allowed us to identify entire regions and individual residues on MscL, which undergo substantial conformational changes upon pocket delipidation (L89W background).

#### Membrane stretching promotes pore hydration in the WT channel

To mimic the naturally occurring activation of MscL by membrane tension, we implemented MD simulations on the WT TbMscL in lipids, with and without tension applied to the lipid bilayer. We performed all simulations with the same lipid composition in order to compare the pore properties and overall channel architecture between the states. Our simulations were initiated from the TbMscL X-ray structure (PDB 2OAR)(55) carrying no modifications. Previously, significantly higher bilayer tension was required to open MscL in MD simulations over short periods of simulation time(37–39), in comparison to the lower tension (~67.5 mN/m) and longer simulation times (300 ns) used in this study, allowing us to capture intermediate states.

WT TbMscL was inserted in a POPC bilayer and simulations for 300 ns were performed in which the x- and y- pressure of the box was set at −50 bar creating constant bilayer tension. During the simulations, the bilayer could expand to adopt to the expansion occurring in the xy plane. As a result, the bilayer thickness decreased approximately 1.2 nm compared to the POPC bilayer without tension (Fig 4B and 5). The channel underwent large conformational changes reaching an overall RMSD of ~14 Å compared to the initial closed structure to adapt to the changes in the bilayer by tilting its TM helices towards the xy plane (Fig 4A, S8A and Video S1). In particular, TM1 and TM2 gradually tilted by 30 and 15 degrees respectively towards the end of the simulations, while the pressure-sensitive pocket surface and the area exposed to the bilayer decreased dramatically (Fig 4, S8B and C). Increasing the timescales of the simulations was outside the scope of this study, as our main interest was to trap and investigate in detail this intermediately open state and the first stop in the MscL’s tension mediated activation pathway. This has enabled us to compare this model of a tension activated state with our experimentally derived state obtained by pocket lipid removal, in the absence of membrane tension (8).

**Figure 4.**
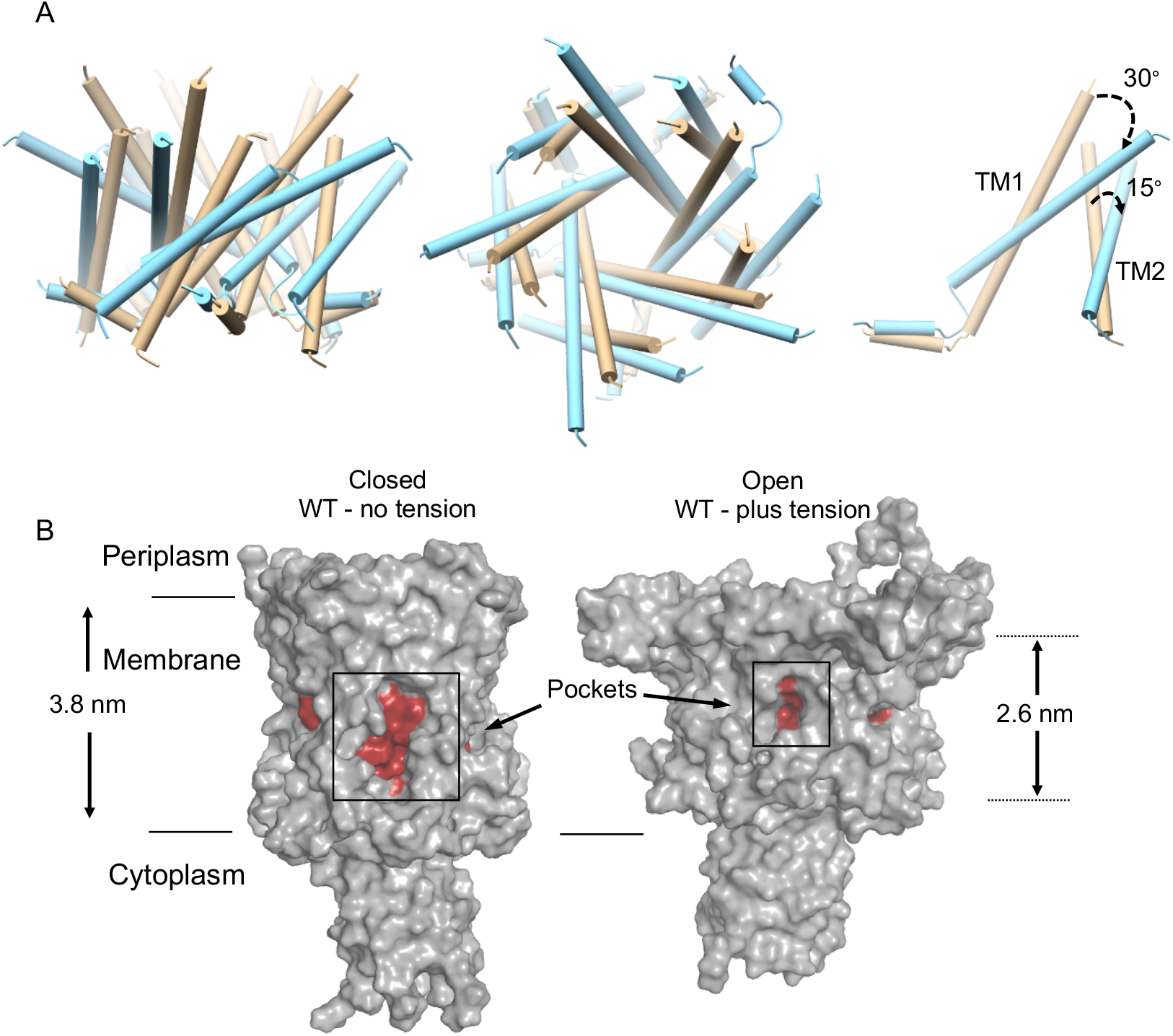
Tension-mediated expanded MscL state. A. TM comparison between closed TbMscL (tan, PDB 2OAR) and the open state WT MscL generated by MD simulations (cyan, under tension), side, top and single subunit views. The latter shows a substantial tilting of TM1 and TM2 towards the membrane plane. These conformational changes occurring under tension are more evident in Video S1. B. The pockets surface dramatically decreases upon tension application, limiting lipid access. Uniform membrane thinning of 1.2 nm (3.8 to 2.6 nm) also occurs during channel expansion, due to tension application.

**Figure 5.**
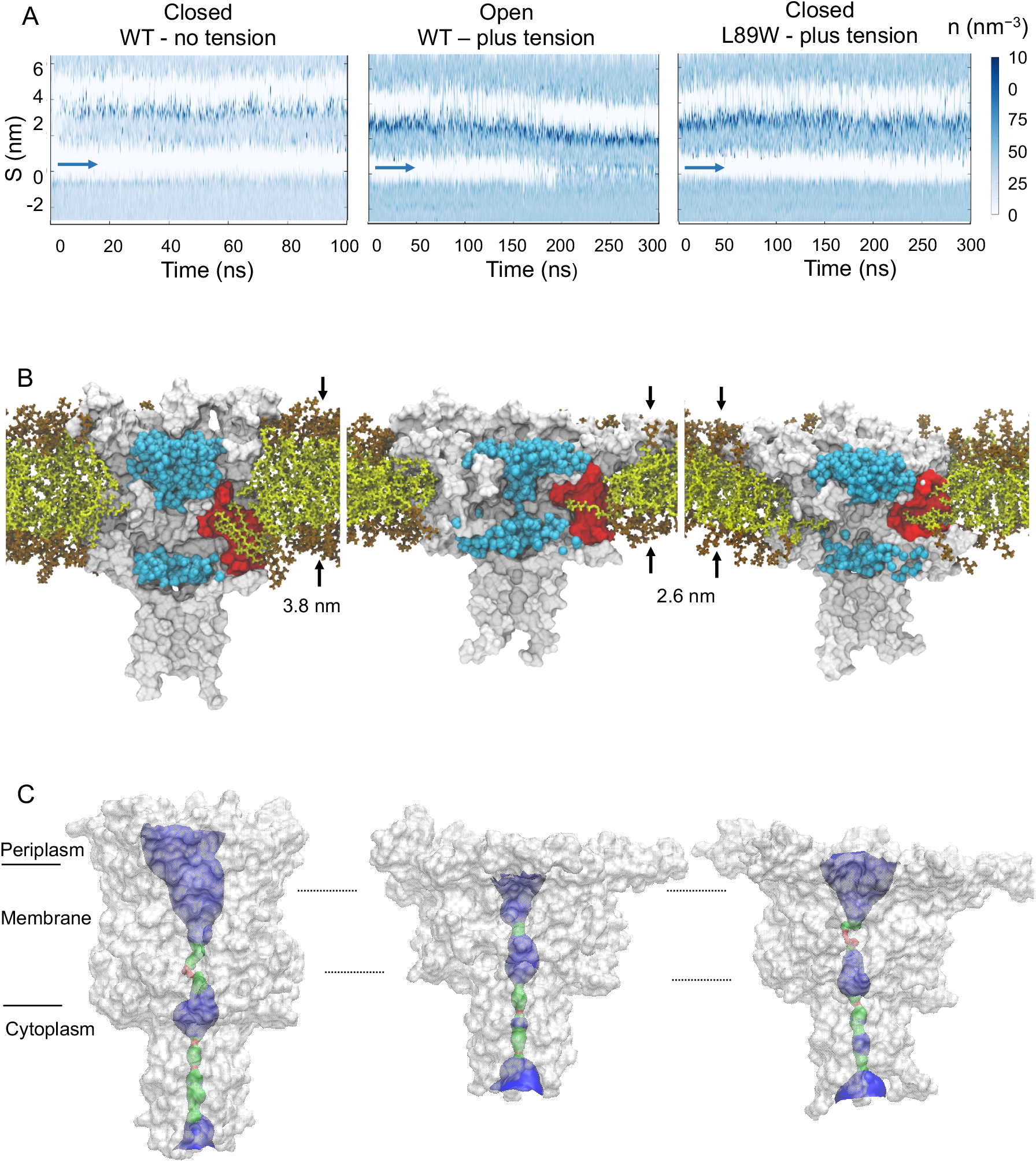
MscL pore hydration investigation under applied bilayer tension. WT under no tension (left column), WT under bilayer tension (right column), L89W under bilayer tension (right column). A. Solvent density profile of the MD simulations using the CHAP(61) software with the vapour-lock position of MscL marked with blue arrows. B. Membrane becomes thinner when tension is applied but only the WT TbMscL pore is hydrated (blue spheres) in contrast to L89W channel. Lipid (olive sticks) availability is larger in the closed state, and lipids have easier access within the pockets to provide force on the back of the vapour lock, keeping the MscL pore closed. This is in contrast to WT MscL under tension (middle column) where the pockets become smaller, while lipid availability decreases, and the pore becomes hydrated. However, when lipids are trapped in the pockets of the L89W modified channel, the pore does not become hydrated despite structural rearrangements occurring (right column). C. Surface visualization of the pore pathway using the program HOLE(62). Red colour indicates a pore radius smaller than 1.15 Å (water molecules cannot go through such an opening), blue represents a radius larger than 2.3 Å and green between 1.15 to 2.3 Å.

Analysis of pore hydration profiles using CHAP(61) showed opening and hydration of the WT channel pore under tension in all of our simulations (Fig 5A). Whilst the higher (compared to biologically relevant) membrane tension does not allow us to provide information about the timescales of the channel transition from the closed to the hydrated state, it enables us to obtain structural data for a key intermediate state, which has a hydrated partially open pore, essential in MscL’s gating process. For these pore hydration events we monitored the annular lipids, which make direct contact or reside in close proximity to the channel and simultaneously interrogated the pore radius profile using HOLE(62) (Fig 5B, C and 6C). For WT, we first observed that following constant tension application for ~200 ns, the pore becomes hydrated, and the channel undergoes major structural rearrangements (Fig 5A). Although a significant membrane thinning (~1.2 nm) occurs, L89W’s pore cannot open to allow water molecules to flow through and the pore radius is still restricted to ~1 Å, despite similar global structure changes as the WT channel (Fig 5 B,C,S8 and Table S2). Following bilayer stretching the total pocket surface area decreases and limits access and availability to bilayer lipids throughout our simulations (Fig 5B). Overall, the under-tension WT MscL MD data agree well with the HDX-MS and 3pESEEM accessibility data obtained for L89W (pocket delipidation by modification). In both cases, similar channel regions become accessible to solvent suggesting a high similarity between the obtained expanded states (Fig 7A).

**Figure 6.**
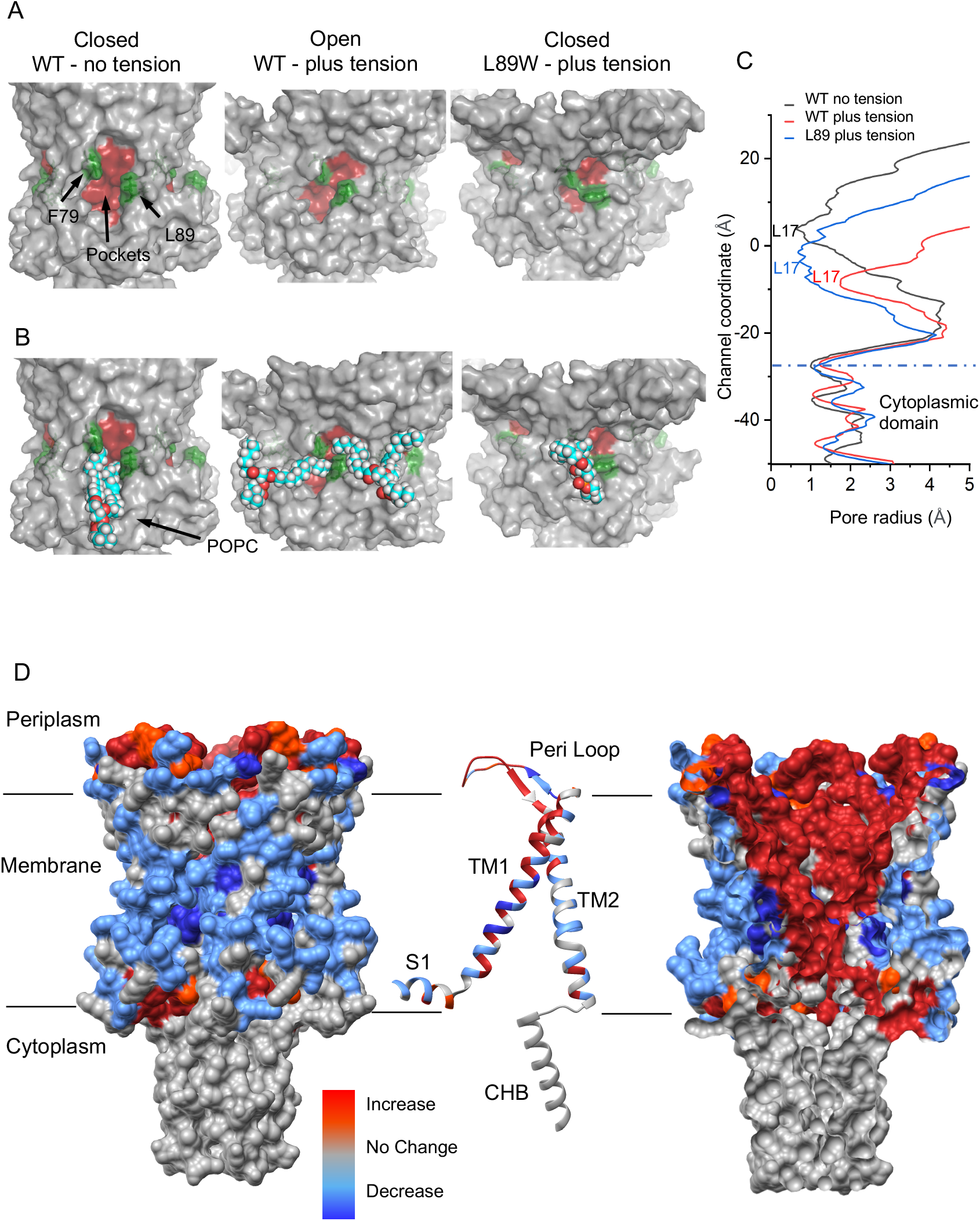
The effect of membrane tension to MscL protein-lipid contact interactions. (A) and (B). F79 and L89W form a “molecular bridge” in the L89W under tension state and consequently trap the lipids within the pockets (right column). F79 and L89 come into greater proximity under bilayer tension in the WT channel (expanded MscL, middle column) compared to the closed state. However, they do not prevent the lipids from exchanging with the bulk bilayer during tension application. The L89W mutation locks lipids in the pockets, preventing the channel from transiting to a hydrated state. C. MscL pore radius profiles analysis and comparisons using HOLE(62). Although L89W MscL undergoes similar major rearrangements to WT MscL under tension its pore remains significantly smaller and almost identical to WT closed MscL under no tension. D. Relative changes in the number of lipid contacts following stimulated tension application in the membrane during MD. The blue regions show decrease in lipid contacts, while the red regions show an increase in lipid contacts. From left to right: Surface view, single subunit cartoon representation and internal MscL surface view. Internal channel regions become more exposed to lipids, while initially membrane exposed regions (closed state) become less exposed to lipids, suggesting a MscL TM helical rotation upon tension application and channel expansion/opening.

**Figure 7.**
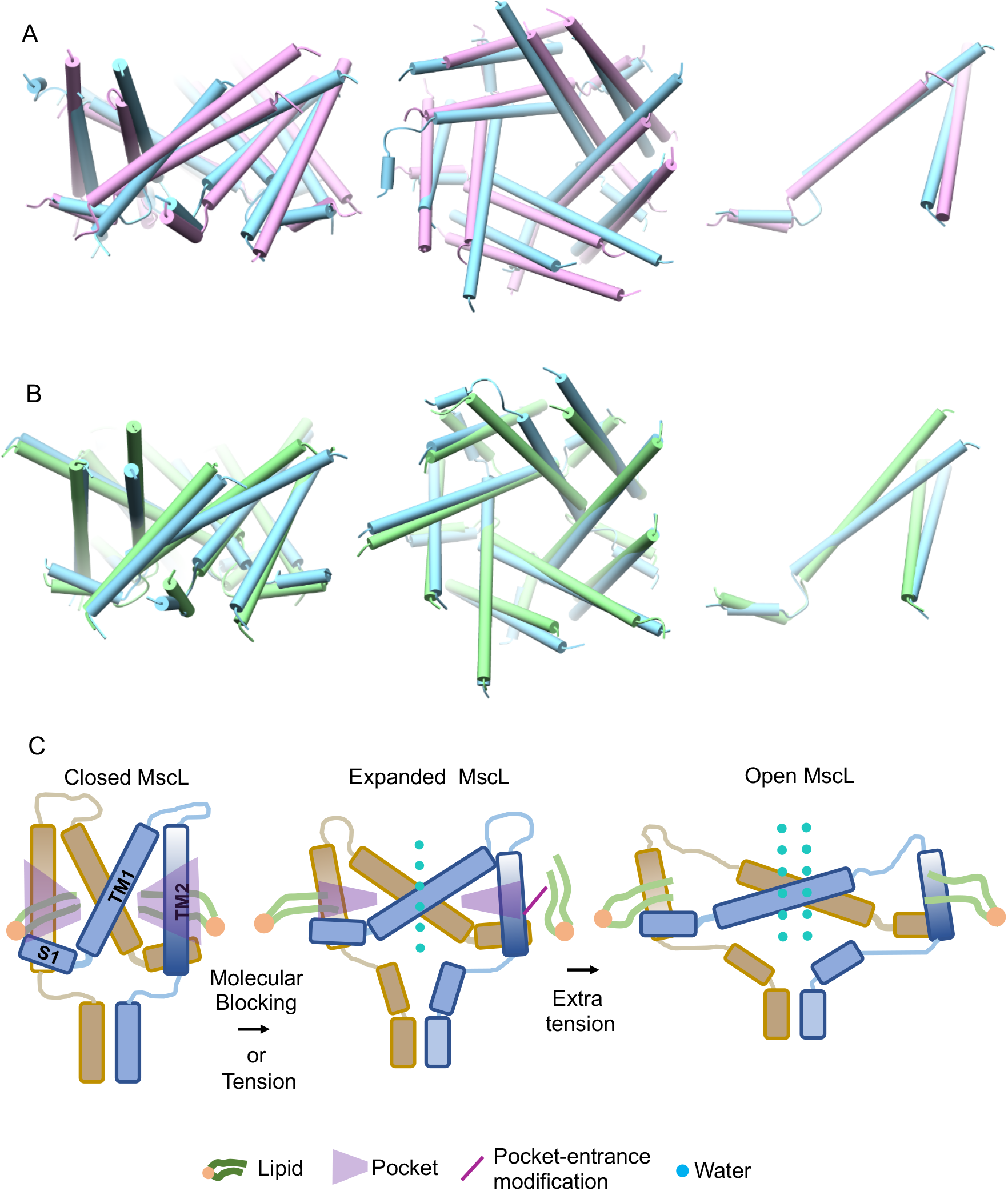
Comparison between tension mediated and pocket delipidation induced MscL states and proposed model. TM1 and TM2 tilt and rotate to initiate MscL opening. Comparison between expanded MaMscL (orchid, PDB ID: 4Y7J) and the open state WT TbMscL generated by MD (cyan, under tension) (A) and the open WT TbMscL state generated by MD (cyan, under tesion) and L89W TbMscL (green, under tension) (B). C. Proposed gating model for pocket lipid removal and tension activated modes. Closed (vertical TM helices, large pockets, tightly associated annular lipids and non-hydrated pore), Expanded (tilted TM helices, smaller pockets, loosely associated annular lipids and hydrated pore). Extra membrane tension is required to fully open MscL.

#### Safety-pin acting trapped lipids in the pockets do not allow MscL pore hydration

A direct consequence of the lipids-move-first model is that if lipids are trapped within the pockets, then the channel pore would not be able to open under membrane tension application. To test this hypothesis, we performed MD simulations with L89W TbMscL under bilayer tension and the same conditions we used for the WT protein (POPC lipid bilayer, pressure value and simulation time). We first tested whether L89W modification causes any distortion on TbMscL’s secondary structure. To this end, we calculated an overall RMSD of 1.8 Å between the L89W and WT TbMscL following equilibration in lipids (and in the absence of applied tension), suggesting there is negligible impact on MscL’s structure imposed by the L89W mutation. We observed that under bilayer tension L89W undergoes similar global structural changes as the WT TbMscL. In all our simulations modified MscL’s pore does not become hydrated, even under the tension conditions which previously promoted WT pore hydration (Fig 5A, 7B, S8 and Table S2).

We observed that during the equilibration of our simulations, lipids intercalated into the pockets and occupied them. Following tension application, lipids were pulled towards the membrane, but the bulky tryptophan modification at the entrance disrupted the lipid exit path, despite tension application sufficient to open the WT channel was applied (Fig 5). Interestingly, in the tension activated state, F79 and L89 from adjacent subunits come very close to form two new smaller pockets in the inner- and outer-leaflets of the now substantially thinner membrane (Fig 6A and B). This spatial “refinement” may facilitate the next stage in MscL’s activation, which follows this sub-conducting state, and requires additional and highly localised pressure for full pore opening of 3 nS and ~30 Å (2, 34, 35). This suggests that the expanded MscL state is an intermediate stop in the tension mediated activation path and not the final destination, e.g. as in activation caused by pocket lipid removal of MscL. Asymmetry in bilayer tension sensitivity has been previously observed for MscL, MscS and other mammalian MS channels (63, 64) and we observed that inner-leaflet lipids previously occupying MscL’s pockets to keep the channel closed, could subsequently penetrate either of the two newly created sub-pockets (Fig 5B, 6A and B). This change restricts the access of new lipids from the bulk membrane to the pockets and immobilizes trapped lipids within the pockets that entered initially during equilibration. In such a case these lipids should interact more strongly with residue 89 in the modified (W89) than the WT (L89) channel. To test this hypothesis, we calculated the pairwise energy forces between all lipids and residue 89, L and W, for the WT and modified channels respectively (Table 1). We found that the lipid interaction (Lennard-Jones) with residue 89 is twice as strong in the modified compared to WT channel (Table 1). This suggests that lipids are “stuck” by the modification in the pockets, reducing their ability to exchange with the bulk bilayer during tension application (Fig 6A and B). Despite the substantial global structural rearrangements, we observed in L89W MscL under tension, when lipids remain tightly associated with the pockets, its pore cannot hydrate (Fig 5 and 6C).

**Table 1.** Pairwise force energy calculations between TbMscL 89 (L or W) and lipids in the WT (L) and pocket-entrance modified (W) channels under tension

| 89 -<br>POPC | Coulomb<br>(KJ/mol) | SD - Coulomb<br>(KJ/mol) | Lennard Jones<br>(KJ/mol) | SD - Lennard<br>Jones (KJ/mol) | Total Energy<br>(KJ/mol) | SD – Total<br>Energy (KJ/mol) |
| --- | --- | --- | --- | --- | --- | --- |
| WT | -14.50 | 1.76 | -101.39 | 27.43 | -115.89 | 29.19 |
| L89W | -14.77 | 3.90 | -192.48 | 3.39 | -207.25 | 7.29 |

#### Lipid order and accessibility during tension application

To assess the lipid contact profile with the MscL TM domain we calculated the relative difference of lipid contacts between the with and no tension TbMscL states (Fig 6 and S9). We found that the relative lipid contacts within bilayer facing residues on the TM domain were reduced over the course of the simulation. In contrast, the lipid contacts of hydrophobic residues that were not lipid exposed in the closed state increased substantially (Fig 6 and S9). This is consistent with TM2 rotation occurring during MscL opening(8) and with pockets decreasing their surface contact area and becoming less accessible to lipids (Fig 4B). To understand lipid rearrangement occurring upon tension application we calculated the lipid order before and after bilayer tension application and found that all lipids (particularly the end of the lipid chains) orient substantially more horizontally in the presence of tension (Fig S10A and C). However, the pocket lipids specifically orient marginally more horizontal compared to the bulk bilayer lipids, in the presence of tension (Fig S10B). These findings suggest that although tension application results in a more horizontal, in respect to the membrane plane, lipid reorientation, additional horizontal tilting other than that caused by membrane stretching is not required to favor lipid pocket preference or specificity (8, 10).

### Discussion

According to the lipids move first model(2), MS channels respond and open their pore when under membrane tension or when lipid occupancy within the pockets is reduced. Equally the channel’s pore will not open, if lipids are trapped within the pockets. It was not previously known whether the functional subconducting intermediate states adopted by MscL upon either membrane tension or pocket delipidation were analogous in MscL. Transitions between functional states of MscL may follow a similar path to cover the available conformational space between these states but may not necessarily sample the exact same discrete stable intermediates. Here we find that both activation mechanisms (Mechanical or pocket lipid removal), result in MscL adopting a well-defined, expanded state.

Our accumulative data suggest this could be the final destination of the pocket delipidation path, but only an intermediate stop of the tension mediated activation path. Therefore, pocket lipid removal constitutes a major component of the initial stages (to open or not to open) of tension-mediated mechanical activation in MscL, consistent with pocket lipids acting as safety-pins of a grenade (8). Following bilayer tension application to WT and L89W channels, we observe similar global structural rearrangements (Fig 5 and 7). Despite this, only the WT channel pore increased in diameter enough for pore hydration. The pore of the L89W channel was significantly smaller in volume and diameter (Fig 2,3,5,6C and S8). Other previously reported loss-of-function or gain-of-unction mutations at, or in proximity to the lipid binding pockets interfere with MscL protein-lipid interactions and may stabilize MscL states by restricting lipid access to the pockets (14, 15). Previously, when lipids were sterically blocked from entering the pockets, the probability of pore opening increased and the activation threshold significantly decreased(8). Equivalently, in our present MD simulations, the L89W mutation locks the exit door and traps the lipids in the pockets by restricting the space available for lipids to move out. Consequently, this disallows channel pore hydration under applied tension, which we indeed observe in our simulations of L89W TbMscL (under membrane tension) (Fig 6A, B and Table 1). Lipids are loosely associated with the pockets in the “hydrated” WT MscL, following tension activation, but these low pairwise force-contacts are not sufficient to close the channel (Fig 5B, 6B and Table 1). We find that the pockets become substantially smaller under tension and consequently less accessible to lipids (Fig 4B, 5B and 6). A similar activation mechanism has been suggested for the small conductance MS channel MscS where inner-leaflet lipids residing within the pockets are required to move before outer-leaflet lipids can disengage to allow gating (25), and the degree of their occupancy within the pockets was linked with distinct MscS states(17, 31). This agrees with our entropy-driven “lipids-move-first model” and with pocket lipids being able to exchange with the bulk bilayer despite its distant positioning away from the pockets (2, 17, 30, 31, 65–67).

In the modified L89W MscL, F79 and W89 form a “molecular” bridge resulting in a two-compartment repartitioning of the smaller pockets (Fig 6A and B). The WT (under tension) expanded state is substantially different from the closed state x-ray structure used (PDB 2OAR) (Fig 4, S8 and Video S1). Importantly, TM2 movement (translation and expansion due to anticlockwise TM2 rotation) under tension like that occurring upon pocket removal activation, consistent with the archaeal x-ray structure of expanded MscL (56), and the expanded (and sub-conducting) state, revealed by PELDOR (2) (Fig 7A).

Previously, cross-sectional expansions not associated with a conducting pore have been proposed for the initial stages of MscL activation(41, 68), suggesting MscL could act as a membrane tension dampener and undergo several transitions with the pore closed (69). Our data support the notion that there are structural transitions that occur prior to channel opening and agree with these models. We observed transitions which are part of the closed-closed expansion gating process, since the overall RMSD between the L89W and WT structures following tension application was small, but they were both significantly different from the closed state (PDB 2OAR) (Fig 4, 7B and S8). This is substantiated by the fact that the channel is unable to reach the intermediate state under applied tension after the lipids are trapped within the pockets *via* the L89W modification, by preventing pore hydration and stabilizing a closed (non-hydrated pore) state (Fig 3, 5, 7).

Our data suggest that for MscL to reach the intermediate hydrated state, triggered either by tension or molecules, the lipids will have to substantially loosen their contacts with the pockets (Fig 5 and Table 1). Lipids could move under tension and adopt a more horizontal orientation in respect to the membrane plane in the WT channel, in the absence of pocket modifications that disrupted lipid exchange with the bulk bilayer (Fig S10). The addition of molecules which could compete for the pockets may support or disrupt lipids acting as negative allosteric modulators within the pocket sites (Fig 7C). Indeed, following similar modifications on MscL, which restrict lipid access to the pockets, or when molecules that specifically bind these pockets (and compete with lipids) were introduced, there was a reduction in MscL’s activation threshold, prolonged openings, increased open channel probability (8, 18, 70).

Disruption of lipid access into the TbMscL’s pockets promoted a mechanical response in the absence of applied tension, while subtle structural differences between these pockets in MscL orthologues dramatically altered their function (8, 19). This pocket region in TbMscL is important for lipid binding (2, 15) and binds specific molecules in EcMscL, which could modulate its activity(16, 18).

Our model is consistent with the notion that other modifications or sites near or within the pocket region could be stabilising MS state(s) by disrupting lipid access to the pockets even more effectively than the L89W modification. Indeed, TbMscL L89 is located at the entrance of the pockets and is structurally aligned to EcMscL M94 and in close proximity to I92, F93, I96 and K97, modifications of which, influence MscL’s activation by ultrasound pulses(71) and decrease its tension threshold for use in a multicompartmental lipid vesicle framework(72). Molecules specific to MscL are expected to act through disruption of lipid-protein interactions(32) within these pockets. These did not fully open the channel, but caused a structural rearrangement of TM2’s cytoplasmic side accompanied by stretching of the cytoplasmic loop(16, 18). This is in agreement with the structural changes observed by HDX and ESEEM for the modified L89W TbMscL channel (or the structurally equivalent M94 EcMscL) and the tension activated WT channel (Fig 1,3,4,7 and S8). V21R1 exposure increases in the intermediate (L89W) MscL state, and this increase is consistent with a channel pore hydration and opening. N13R1’s side chain is already solvent exposed in the closed state, as it is pointing into the cytoplasm (Fig S7), and thus unaffected in the expanded intermediate state (Fig 3).

Transition to the intermediate state increased solvent accessibility on the upper TM1 and TM2, the cytoplasmic loops and the top portion of the cytoplasmic helical bundle (Fig 1 and 2), consistent with stretching of the cytoplasmic loops, of which shortening influences MscL’s gating properties, pore conductance and oligomeric assembly(16, 42, 60, 73).

The top of TM1 (residues 36 - 41) partially unfolds within a single subunit of the MscL pentamer, while the remaining four subunits bend without fully converting into a loop. This finding was consistently observed across all three MD simulation repeats following the symmetric application of tension along the membrane xy plane and suggests the presence of a subunit asymmetry during MscL opening(41, 74, 75). In contrast to the WT channel, all five TM1 helices bend without a break in helical symmetry or conversion into a loop in the L89W channel (Fig 4). For the top of the TM1 helix to partially unfold, lipids should become loosely associated with the pockets before asymmetric opening could occur and eventually lead to full channel opening. Whether the lipids are sequentially released (or loosen their contact) following single subunit disengagement remains to be elucidated. Shifting of the equilibrium occurs in order to destabilize the closed state and initiate symmetric (simultaneous) or asymmetric (sequential) pocket lipid removal from the five MscL subunits. Our data are compatible with such an asymmetry in opening and this structural effect may be the consequence of lipids moving out from MscL’s five pockets sequentially(75). Subsequently, following initial expansion and conversion into loops, these loops could then provide membrane handles for MscL in order to transit to its full opening state (Video S1).

Despite a significant membrane thinning (~1.2 nm), L89W’s pore does not allow water molecules to flow through (Fig 4 and 5). Membrane thickness and hydrophobic mismatch energetically contribute to MscL’s gating, but our data suggest they are not the sole driving force and pocket lipids have to be released first in order to destabilize MscL’s closed state. However, it should be noted that while tension application and channel gating occur within the 300 ns that our simulations run, this may be faster compared to the timescale of MscL opening transitions *in situ*.

Lipids must adopt certain angles in respect to the membrane plane in order to enter (or exit) the pockets and these angles should be consistent with lipids oriented horizontally, contrary to the bulk bilayer lipids. Such lipids have been resolved in multiple x-ray and cryoEM structures of MscS(2, 17, 30), YnaI(28), MSL1(29), TRAAK(26), TREK-2(27) and TRPV3(25) channels. When we calculated the lipid order parameters for our MD simulations, we found that under tension application the lipids adopt a substantially more horizontal orientation (Fig S10). This is consistent with significantly lower pulling forces required to remove single lipids from TbMscL’s pockets when they are applied at 45o in respect to the membrane plane(76).

In conclusion, here we have generated tension- and modification- activated MscL states and interrogated their pore hydration properties, lipid-protein interactions and monitored their structural dynamics by experimental (HDX-MS and 3pESEEM solvent accessibility measurements) and computational (MD simulations) tools. Accumulatively, our data demonstrate that the intermediate expanded MscL state derived by pocket delipidation (L89W) is structurally analogous to the tension activated MscL state (mechanical)(Fig 7C). Our data further suggest that the expanded MscL state is the final destination of MscL’s pocket lipid removal activation path, but only an intermediate stop for the tension mediated activation path, which leads to a fully open MscL state. It is quite intriguing that whether it is by tension application or pocket lipid depletion, we find that MscL is sampling similar defined states, as is also the case for MscS, whose states are also guided by pocket-lipid availability (17).

The structural similarities of these two differently derived states suggest that lipids could act as molecular triggers on specific sites and mimic the naturally occurring tension activation in MS channels, with implications for MS channel evolution and multimodality.

### Materials and Methods

#### Materials

N-Dodecyl-b-D-Maltopyranoside (DDM) anagrade was obtained from Anatrace or Glycon (Germany). Isopropyl-β-D-thiogalactoside (IPTG) was obtained from Formedium, and (Tris(2-carboxyethyl)phosphine (TCEP) was obtained from Thermo Scientific. The S-(2,2,5,5-tetramethyl-2,5-dihydro-1H-pyrrol-3-yl)methyl methanesulfonothioate (MTSSL) spin label was obtained from Toronto Research Chemicals. Lipids were purchased from Avanti Polar Lipids and Biobeads from Biorad. All other chemicals unless otherwise stated were obtained from Sigma.

#### Methods

##### Mutagenesis and Protein Expression

Mutagenesis of TbMscL with a C-terminal 6xHis-tag in a pJ411:140126 vector allowed the generation of single and double mutants. Primers were used for site-directed mutagenesis reactions to introduce the point mutations. The wild type and mutant plasmids were transformed into BL21(DE3) (ThermoFisher) E. coli cells. Cells were grown in 550 ml LB medium in a 2L flask at 37 °C until they reached an OD600 ~0.8 and subsequently cooled down to 25 °C and induced with 1 mM IPTG for 3.5 hrs. Cells were harvested by centrifugation at 4000 × g and stored at - 80 °C, until further use.

#### Protein Purification and Spin Labelling

The protocols followed in this study were similar to the ones previously described(8, 19, 35). In brief, following protein expression, cell pellets were resuspended in phosphate-buffered saline and subjected to lysis using a cell disrupter at 30 kpsi. To remove cell debris the suspension was centrifuged at 4000 × g for 20 min and the resulting supernatant was centrifuged again at 100,000 × g for 1 h. The membrane pellet was resuspended and solubilized in buffer containing 50 mM sodium phosphate at pH 7.5, 300 mM NaCl, 10% v/v glycerol, 50 mM imidazole and 1.5% w/v DDM (solubilization buffer) and incubated at 4 °C for 1 h. The sample was then centrifuged at 4000 × g for 20 min and the supernatant was passed through a +2Ni-NTA column containing 0.75 mL of +2Ni-NTA beads. The column was then washed with 10 mL buffer containing 50 mM sodium phosphate of pH 7.5, 300 mM NaCl, 10% v/v glycerol and 0.05% w/v DDM (wash buffer) and then with 5 mL wash buffer supplemented with 3 mM TCEP, for reduction of the cysteines. Afterwards, MTSSL dissolved in wash buffer at a 10-fold excess of the expected protein concentration was added to the column and left to react for 2 h at 4 °C. The protein was then eluted from the column with 5 mL of wash buffer supplemented with 300 mM imidazole. Finally, the protein was subjected to SEC using a Superdex 200 column (GE Healthcare) equilibrated with buffer containing 50 mM sodium phosphate of pH 7.5, 300 mM NaCl and 0.05% w/v DDM. Collected fractions of TbMscL were then concentrated to ~800 μM monomer concentration, which is suitable for the EPR samples preparation

##### 3pESEEM sample preparation

Purified TbMscL detergent or reconstituted samples were diluted by 50% with deuterated ethylene glycol as cryoprotectant to a final monomer concentration of ~400 μM and a volume of 70 μL of the mixture was then transferred into a 3 mm (OD) quartz tubes and flash frozen in liquid N2.

##### 3pESEEM spectroscopy

X-band three pulse ESEEM measurements were performed on a Bruker ELEXSYS E580 spectrometer with a 4 mm dielectric resonator (MD4). Measurements were conducted at 80 K. The 3-pulse sequence used for the experiments is π/2-τ-π/2-T-π/2-stimulated echo with a pulse length t_π/2_ = 16 ns and inter-pulse delay τ that was adjusted to match either the blind spots of the proton or deuterium. The delay T was incremented from 400ns in 12 ns steps. A four-step phase cycling was used to eliminate the unwanted echoes. All the measurements were performed at the maximum of the filed sweep spectrum of the nitroxide. For solvent accessibility determination, only-3pESEEM data recorded with τ that corresponds to the proton blind spot were used. The obtained time-domain traces were background-corrected, apodized with a hamming window and zero-filled prior to Fourier transformation. The solvent accessibility can be determined by different analysis methods from the deuterium 3pESEEM. In the present study, we used a model developed by Jeschke and co-workers and it is based on deuterium modulation depth(51). The primary ESEEM data was background corrected in a such way to obtain a deconvoluted and normalized nuclear modulation function. We used two approaches to determine the solvent accessibility; the first one by fitting the obtained nuclear modulation function for each mutant by a damped harmonic oscillation function and the outcome from the fitting was used to estimate the solvent accessibility(51), the second approach by Fourier transformation of the nuclear modulation function and in this case the water accessibility was determined directly form the intensities of the deuterium peaks in the magnitude ESEEM spectra. The solvent accessibility parameters derived from both methods are in a good agreement within the error bars as shown in Fig S3. Although the standard deviations for the fitted parameters were less than 2% for all the mutants, we additionally accounted for errors that might emanate from differences in relaxation behavior or in contribution from no-water protons between the different mutants and use an error of 5%. All ESEEM raw data for this study are within the following link: http://archive.researchdata.leeds.ac.uk/777/

##### HDX-MS sample preparation

For HDX WT or L89W mutant MscL were not spin-labelled. The cell disruption, membrane preparation, solubilization, and application of the solubilized membranes to the Ni-NTA column occurred as previously. The column was washed, as previously, with 10 mL buffer containing 50 mM sodium phosphate of pH 7.5, 300 mM NaCl, 10% v/v glycerol and 0.05% w/v DDM (wash buffer). Following this step, the protein was eluted with 5 ml of wash buffer containing 300 mM imidazole. The protein was not treated with TCEP or MTSL supplemented wash buffer as the protein was not spin-labelled. The protein was then subjected to SEC using a Superdex 200 column as previously described. The fractions were collected, and the protein samples were concentrated to achieve 350 μl of protein at 16 μM.

##### HDX and LC-MS/MS

HDX-MS experiments were carried out using an automated HDX robot (LEAP Technologies, Fort Lauderdale, FL, USA) coupled to an M-Class Acquity LC and HDX manager (Waters Ltd., Wilmslow, Manchester, UK). All samples were diluted to 16 μM in equilibration buffer (50 mM potassium phosphate, 300 mM NaCl, pH 7.4, 0.05% DDM) prior to analysis. 5 μL sample was added to 95 μl deuterated buffer (50 mM potassium phosphate, 300 mM NaCl pD 7.4, 0.05% DDM) and incubated at 4 °C for 0.5, 1, 2, 10 or 60 min. Following the labelling reaction, samples were quenched by adding 50 μl of the labelled solution to 100 μl quench buffer (50 mM potassium phosphate, 300 mM NaCl, 0.1% DDM pH 2.2) giving a final quench pH ~2.5. 50 μl of quenched sample was injected on to the HDX Manger and passed through immobilized pepsin and aspergillopepsin columns connected in series (AffiPro, Czech Republic) at 115 μl min-1 (20 °C) and loaded on to a VanGuard Pre-column Acquity UPLC BEH C18 (1.7 μm, 2.1 mm X 5 mm, Waters Ltd., Wilmslow, Manchester, UK) for 3 mins in 0.3 % formic acid in water. The resulting peptides were transferred to a C18 column (75 μm X 150 mm, Waters Ltd., Wilmslow, Manchester, UK) and separated by gradient elution of 0-40 % MeCN (0.1 % v/v formic acid) in H2O (0.3 % v/v formic acid) over 7 mins at 40 μl.min-1. Trapping and gradient elution of peptides was performed at 0 oC to reduce back exchange. The HDX system was interfaced to a Synapt G2Si mass spectrometer (Waters Ltd., Wilmslow, Manchester, UK). HDMSE and dynamic range extension modes (Data Independent Analysis (DIA) coupled with IMS separation) were used to separate peptides prior to CID fragmentation in the transfer cell (77). HDX data were analyzed using PLGS (v3.0.2) and DynamX (v3.0.0) software supplied with the mass spectrometer. Restrictions for identified peptides in DynamX were as follows: minimum intensity: 1000, minimum products per MS/MS spectrum: 5, minimum products per amino acid: 0.3, maximum sequence length: 25, maximum ppm error: 5, file threshold: 3/3. Following manual curation of the data, PAVED(78) and Deuteros(79) were used to identify peptides with statistically significant increases/decreases in deuterium uptake (applying 99% or 95% confidence intervals) and to prepare Wood’s plots. The raw HDX-MS data, have been deposited to the ProteomeXchange Consortium via the PRIDE partner repository with the dataset identifier PXD021983. A summary of the HDX-MS data, as recommended by reporting guidelines, is shown in Table S1. All data is available via ProteomeXchange with identifier PXD021983. Project Name: Mechanical and molecular activation lead to structurally analogous MscL states. Project accession: PXD021983 Username, Password: lnAkk2H3

##### Atomistic molecular dynamics simulations under no tension

The set up and parameters of MD simulations without tension were as previously described(8). In brief, CHARMM-GUI was used to insert the TbMscL structure (2OAR) into a pre-equilibrated patch of POPC bilayer containing approximately 387 lipids and occupying an area of 120 × 120 Å2. The protein and membrane bilayer were solvated with TIP3P water and 150 mM NaCl. The simulations were performed in an NPT ensemble at 303.15 K and 1bar pressure on all xyz axes using GROMACS_2016.4 with CHARMM36 force field. The particle mesh Ewald (PME) method was applied to calculate electrostatic forces, and the van der Waals interactions were smoothly switched off at 10–12 Å by the force-switch manner. The time step was set to 2 fs in conjunction with the LINCS algorithm. After the standard minimization and equilibration steps using the GROMACS input scripts generated by CHARMM-GUI, 100 ns dynamic simulation was calculated.

#### Atomistic molecular dynamics simulations under bilayer tension

These MD simulations were also set up using CHARMM-GUI(80). The MscL structure (PDB 2OAR) in the wild type and in a mutated form (L89W) was inserted in a symmetric bilayer containing 514 POPC lipids. Systems were neutralized with a 150 mM concentration of NaCl. The obtained system was energy minimised for 5000 steps and subsequently equilibrated in 6 steps following CHARMM-GUIs equilibration. Stretch-induced conformational changes in both the wild-type and mutant MscL were investigated by unrestrained simulations of 300 ns (3 simulation repeats for the WT and 2 for the L89W channel) where the bilayer plane (xy plane) pressure was changed semiisotropically to - 50 bars (which corresponds to a tension of ~67.5 mN/m). The pressure in the bilayer normal (z direction) was kept at +1 bar and the temperature at 310 K. All the atomistic systems were simulated using GROMACS 2016(81) with CHARMM36 force field(82) and a 2 fs time step. The Nose-Hoover thermostat(83) and the Parrinello-Rahman barostat(84) were used for the stretch-induced simulations. Long-range electrostatics were managed using the particle-mesh Ewald method(85) and the LINCS algorithm was used to constrain bond lengths(86). All MD trajectories and data for this study is within the following link: http://archive.researchdata.leeds.ac.uk/777/

## Supporting information

Suppl_Tables and Figures_Wang_Lane et al_Pliotas_2021

## Data accessibility

HDX mass spectrometry:

Data are available via ProteomeXchange with identifier PXD021983.
Project Name: Mechanical and molecular activation lead to structurally analogous MscL states
Project accession: PXD021983
Reviewer account details: Username, Password: lnAkk2H3

3pESEEM and MD:

Data are available within the following link: http://archive.researchdata.leeds.ac.uk/777/

## Acknowledgments

This project was supported by a Biotechnology and Biological Sciences Research Council (BBSRC) grant (BB/S018069/1) to C.P., who also acknowledges support from the Wellcome Trust (WT) (219999/Z/19/Z) and the Chinese Scholarship Council (CSC) in the form of studentships for B.J.L. and B.W. respectively. ANC is a Sir Henry Dale Fellow jointly funded by the WT and the Royal Society (220628/Z/20/Z). Funding from the BBSRC (BB/M012573/1) enabled the purchase of mass spectrometry equipment.

## References

1. Pliotas C & Naismith JH (2016) Spectator no more, the role of the membrane in regulating ion channel function. Curr Opin Struct Biol 45:59–66.

2. Pliotas C, et al. (2015) The role of lipids in mechanosensation. Nat Struct Mol Biol 22(12):991–998.

3. Teng J, Loukin S, Anishkin A, & Kung C (2015) The force-from-lipid (FFL) principle of mechanosensitivity, at large and in elements. Pflugers Arch 467(1):27–37.

4. Malcolm HR, Blount P, & Maurer JA (2015) The mechanosensitive channel of small conductance (MscS) functions as a Jack-in-the box. Biochim Biophys Acta 1848(1 Pt A):159–166.

5. Cox CD, Nakayama Y, Nomura T, & Martinac B (2015) The evolutionary ‘tinkering’ of MscS-like channels: generation of structural and functional diversity. Pflugers Arch 467(1):3–13.

6. Brohawn SG (2015) How ion channels sense mechanical force: insights from mechanosensitive K2P channels TRAAK, TREK1, and TREK2. Ann N Y Acad Sci 1352:20–32.

7. Naismith JH & Booth IR (2012) Bacterial mechanosensitive channels--MscS: evolution’s solution to creating sensitivity in function. Annu Rev Biophys 41:157–177.

8. Kapsalis C, et al. (2019) Allosteric activation of an ion channel triggered by modification of mechanosensitive nano-pockets. Nat Commun 10(1):4619.

9. Patrick JW, et al. (2018) Allostery revealed within lipid binding events to membrane proteins. Proc Natl Acad Sci U S A 115(12):2976–2981.

10. Laganowsky A, et al. (2014) Membrane proteins bind lipids selectively to modulate their structure and function. Nature 510(7503):172–175.

11. Ridone P, et al. (2018) “Force-from-lipids” gating of mechanosensitive channels modulated by PUFAs. J Mech Behav Biomed Mater 79:158–167.

12. Ward R, et al. (2014) Probing the structure of the mechanosensitive channel of small conductance in lipid bilayers with pulsed electron-electron double resonance. Biophys J 106(4):834–842.

13. Pliotas C, et al. (2012) Conformational state of the MscS mechanosensitive channel in solution revealed by pulsed electron-electron double resonance (PELDOR) spectroscopy. Proc Natl Acad Sci U S A 109(40):E2675–2682.

14. Anishkin A, Chiang CS, & Sukharev S (2005) Gain-of-function mutations reveal expanded intermediate states and a sequential action of two gates in MscL. J Gen Physiol 125(2):155–170.

15. Iscla I, Wray R, Eaton C, & Blount P (2015) Scanning MscL Channels with Targeted Post-Translational Modifications for Functional Alterations. PLoS One 10(9):e0137994.

16. Wray R, et al. (2016) Dihydrostreptomycin Directly Binds to, Modulates, and Passes through the MscL Channel Pore. PLoS biology 14(6):e1002473.

17. Zhang Y, et al. (2021) Visualization of the mechanosensitive ion channel MscS under membrane tension. Nature 590:509–514.

18. Wray R, Iscla I, Kovacs Z, Wang J, & Blount P (2019) Novel compounds that specifically bind and modulate MscL: insights into channel gating mechanisms. FASEB J 33(3):3180–3189.

19. Kapsalis C, Ma Y, Bode BE, & Pliotas C (2020) In-Lipid Structure of Pressure-Sensitive Domains Hints Mechanosensitive Channel Functional Diversity. Biophys J 119(2):448–459.

20. Syeda R, et al. (2015) Chemical activation of the mechanotransduction channel Piezo1. Elife 4.

21. Botello-Smith WM, et al. (2019) A mechanism for the activation of the mechanosensitive Piezo1 channel by the small molecule Yoda1. Nat Commun 10(1):4503.

22. Bagriantsev SN, Peyronnet R, Clark KA, Honore E, & Minor DL, Jr. (2011) Multiple modalities converge on a common gate to control K2P channel function. EMBO J 30(17):3594–3606.

23. Piechotta PL, et al. (2011) The pore structure and gating mechanism of K2P channels. EMBO J 30(17):3607–3619.

24. Feliciangeli S, Chatelain FC, Bichet D, & Lesage F (2015) The family of K2P channels: salient structural and functional properties. J Physiol 593(Pt 12):2587–2603.

25. Deng Z, et al. (2020) Gating of human TRPV3 in a lipid bilayer. Nat Struct Mol Biol.

26. Brohawn SG, Campbell EB, & MacKinnon R (2014) Physical mechanism for gating and mechanosensitivity of the human TRAAK K+ channel. Nature 516(7529):126–130.

27. Dong YY, et al. (2015) K2P channel gating mechanisms revealed by structures of TREK-2 and a complex with Prozac. Science 347(6227):1256–1259.

28. Flegler VJ, et al. (2020) The MscS-like channel YnaI has a gating mechanism based on flexible pore helices. Proc Natl Acad Sci U S A 117(46):28754–28762.

29. Deng Z, et al. (2020) Structural mechanism for gating of a eukaryotic mechanosensitive channel of small conductance. Nat Commun 11(1):3690.

30. Rasmussen T, Flegler VJ, Rasmussen A, & Bottcher B (2019) Structure of the Mechanosensitive Channel MscS Embedded in the Membrane Bilayer. J Mol Biol.

31. Flegler VJ, et al. (2021) Mechanosensitive channel gating by delipidation. Proc Natl Acad Sci U S A 118(33).

32. Bavi N, et al. (2016) The role of MscL amphipathic N terminus indicates a blueprint for bilayer-mediated gating of mechanosensitive channels. Nat Commun 7:11984.

33. Erdogmus S, et al. (2019) Helix 8 is the essential structural motif of mechanosensitive GPCRs. Nat Commun 10(1):5784.

34. Hartley AM, Ma Y, Lane BJ, Wang B, & Pliotas C (2021) Using pulsed EPR in the structural analysis of integral membrane proteins. Electron Paramag Reson 27:74–108.

35. Pliotas C (2017) Ion Channel Conformation and Oligomerization Assessment by Site-Directed Spin Labeling and Pulsed-EPR. Methods in enzymology 594:203–242.

36. Debruycker V, et al. (2020) An embedded lipid in the multidrug transporter LmrP suggests a mechanism for polyspecificity. Nat Struct Mol Biol 27(9):829–835.

37. Martinac AD, Bavi N, Bavi O, & Martinac B (2017) Pulling MscL open via N-terminal and TM1 helices: A computational study towards engineering an MscL nanovalve. PLoS One 12(8):e0183822.

38. Gullingsrud J & Schulten K (2003) Gating of MscL studied by steered molecular dynamics. Biophys J 85(4):2087–2099.

39. Bavi N, et al. (2017) Nanomechanical properties of MscL alpha helices: A steered molecular dynamics study. Channels 11(3):209–223.

40. Jeon J & Voth GA (2008) Gating of the mechanosensitive channel protein MscL: the interplay of membrane and protein. Biophys J 94(9):3497–3511.

41. Konijnenberg A, et al. (2014) Global structural changes of an ion channel during its gating are followed by ion mobility mass spectrometry. Proc Natl Acad Sci U S A 111(48):17170–17175.

42. Herrera N, Maksaev G, Haswell ES, & Rees DC (2018) Elucidating a role for the cytoplasmic domain in the Mycobacterium tuberculosis mechanosensitive channel of large conductance. Sci Rep 8(1):14566.

43. Mukherjee N, et al. (2014) The activation mode of the mechanosensitive ion channel, MscL, by lysophosphatidylcholine differs from tension-induced gating. FASEB J 28(10):4292–4302.

44. Katsuta H, Sawada Y, & Sokabe M (2019) Biophysical Mechanisms of Membrane-Thickness-Dependent MscL Gating: An All-Atom Molecular Dynamics Study. Langmuir 35(23):7432–7442.

45. Aryal P, et al. (2017) Bilayer-Mediated Structural Transitions Control Mechanosensitivity of the TREK-2 K2P Channel. Structure 25(5):708–718.e702.

46. Martens C, Shekhar M, Lau AM, Tajkhorshid E, & Politis A (2019) Integrating hydrogen-deuterium exchange mass spectrometry with molecular dynamics simulations to probe lipid-modulated conformational changes in membrane proteins. Nat Protoc 14(11):3183–3204.

47. Ahdash Z, et al. (2019) HDX-MS reveals nucleotide-dependent, anti-correlated opening and closure of SecA and SecY channels of the bacterial translocon. Elife 8.

48. Konermann L, Pan J, & Liu YH (2011) Hydrogen exchange mass spectrometry for studying protein structure and dynamics. Chem Soc Rev 40(3):1224–1234.

49. Moller IR, et al. (2019) Conformational dynamics of the human serotonin transporter during substrate and drug binding. Nat Commun 10(1):1687.

50. Cieslak JA, Focia PJ, & Gross A (2010) Electron spin-echo envelope modulation (ESEEM) reveals water and phosphate interactions with the KcsA potassium channel. Biochemistry 49(7):1486–1494.

51. Volkov A, Dockter C, Bund T, Paulsen H, & Jeschke G (2009) Pulsed EPR determination of water accessibility to spin-labeled amino acid residues in LHCIIb. Biophys J 96(3):1124–1141.

52. Liu L, et al. (2016) Probing the Local Secondary Structure of Human Vimentin with Electron Spin Echo Envelope Modulation (ESEEM) Spectroscopy. J Phys Chem B 120(48):12321–12326.

53. Zhang R, et al. (2015) Development of electron spin echo envelope modulation spectroscopy to probe the secondary structure of recombinant membrane proteins in a lipid bilayer. Protein Sci 24(11):1707–1713.

54. Matalon E, et al. (2013) Topology of the trans-membrane peptide WALP23 in model membranes under negative mismatch conditions. J Phys Chem B 117(8):2280–2293.

55. Chang G, Spencer RH, Lee AT, Barclay MT, & Rees DC (1998) Structure of the MscL homolog from Mycobacterium tuberculosis: a gated mechanosensitive ion channel. Science 282(5397):2220–2226.

56. Li J, et al. (2015) Mechanical coupling of the multiple structural elements of the large-conductance mechanosensitive channel during expansion. Proc Natl Acad Sci U S A 112(34):10726–10731.

57. Powl AM, East JM, & Lee AG (2005) Heterogeneity in the binding of lipid molecules to the surface of a membrane protein: Hot spots for anionic lipids on the mechanosensitive channel of large conductance MscL and effects on conformation. Biochemistry 44(15):5873–5883.

58. Michou M, Kapsalis C, Pliotas C, & Skretas G (2019) Optimization of Recombinant Membrane Protein Production in the Engineered Escherichia coli Strains SuptoxD and SuptoxR. ACS Synth Biol 8(7):1631–1641.

59. Bavi N, et al. (2017) Structural Dynamics of the MscL C-terminal Domain. Sci Rep 7(1):17229.

60. Yang LM, et al. (2012) Three routes to modulate the pore size of the MscL channel/nanovalve. ACS Nano 6(2):1134–1141.

61. Klesse G, Rao S, Sansom MSP, & Tucker SJ (2019) CHAP: A Versatile Tool for the Structural and Functional Annotation of Ion Channel Pores. J Mol Biol 431(17):3353–3365.

62. Smart OS, Goodfellow JM, & Wallace BA (1993) The pore dimensions of gramicidin A. Biophys J 65(6):2455–2460.

63. Perozo E, Kloda A, Cortes DM, & Martinac B (2002) Physical principles underlying the transduction of bilayer deformation forces during mechanosensitive channel gating. Nat Struct Biol 9(9):696–703.

64. Belyy V, Kamaraju K, Akitake B, Anishkin A, & Sukharev S (2010) Adaptive behavior of bacterial mechanosensitive channels is coupled to membrane mechanics. J Gen Physiol 135(6):641–652.

65. Anishkin A, Akitake B, & Sukharev S (2008) Characterization of the resting MscS: modeling and analysis of the closed bacterial mechanosensitive channel of small conductance. Biophys J 94(4):1252–1266.

66. Reddy B, Bavi N, Lu A, Park Y, & Perozo E (2019) Molecular basis of force-from-lipids gating in the mechanosensitive channel MscS. Elife 8.

67. Kefauver JM, Ward AB, & Patapoutian A (2020) Discoveries in structure and physiology of mechanically activated ion channels. Nature 587(7835):567–576.

68. Betanzos M, Chiang CS, Guy HR, & Sukharev S (2002) A large iris-like expansion of a mechanosensitive channel protein induced by membrane tension. Nat Struct Biol 9(9):704–710.

69. Boucher PA, Morris CE, & Joos B (2009) Mechanosensitive closed-closed transitions in large membrane proteins: osmoprotection and tension damping. Biophys J 97(10):2761–2770.

70. Iscla I, Wray R, Wei S, Posner B, & Blount P (2014) Streptomycin potency is dependent on MscL channel expression. Nat Commun 5:4891.

71. Ye J, et al. (2018) Ultrasonic Control of Neural Activity through Activation of the Mechanosensitive Channel MscL. Nano Lett 18(7):4148–4155.

72. Hindley JW, et al. (2019) Building a synthetic mechanosensitive signaling pathway in compartmentalized artificial cells. Proc Natl Acad Sci U S A 116(34):16711–16716.

73. Reading E, et al. (2015) The Effect of Detergent, Temperature, and Lipid on the Oligomeric State of MscL Constructs: Insights from Mass Spectrometry. Chem Biol 22(5):593–603.

74. Birkner JP, Poolman B, & Kocer A (2012) Hydrophobic gating of mechanosensitive channel of large conductance evidenced by single-subunit resolution. Proc Natl Acad Sci U S A 109(32):12944–12949.

75. Mika JT, Birkner JP, Poolman B, & Kocer A (2013) On the role of individual subunits in MscL gating: “all for one, one for all?”. FASEB J 27(3):882–892.

76. Vanegas JM & Arroyo M (2014) Force transduction and lipid binding in MscL: a continuum-molecular approach. PLoS One 9(12):e113947.

77. Cryar A, Groves K, & Quaglia M (2017) Online Hydrogen-Deuterium Exchange Traveling Wave Ion Mobility Mass Spectrometry (HDX-IM-MS): a Systematic Evaluation. J Am Soc Mass Spectrom 28(6):1192–1202.

78. Cornwell O, Radford SE, Ashcroft AE, & Ault JR (2018) Comparing Hydrogen Deuterium Exchange and Fast Photochemical Oxidation of Proteins: a Structural Characterisation of Wild-Type and DeltaN6 beta2-Microglobulin. J Am Soc Mass Spectrom 29(12):2413–2426.

79. Lau AMC, Ahdash Z, Martens C, & Politis A (2019) Deuteros: software for rapid analysis and visualization of data from differential hydrogen deuterium exchange-mass spectrometry. Bioinformatics 35(17):3171–3173.

80. Jo S, Kim T, Iyer VG, & Im W (2008) CHARMM-GUI: a web-based graphical user interface for CHARMM. J Comput Chem 29(11):1859–1865.

81. Abraham MJ, T. Murtola, R. Schulz, S. Páll, J.C. Smith, B. Hess, E. Lindah, and E. Lindahl (2015) GROMACS: High performance molecular simulations through multi-level parallelism from laptops to supercomputers. SoftwareX 1–2:19–25.

82. Lee S, et al. (2014) CHARMM36 united atom chain model for lipids and surfactants. J Phys Chem B 118(2):547–556.

83. Evans DJ, Holian B.L. (1985) The Nose-Hoover thermostat. J Chem Phys 83:4069–4074.

84. Parrinello M, Rahman A. (1981) Polymorphic transitions in single crystals: A new molecular dynamics method. J. Appl. Phys. 52:7182–7190.

85. Darden T, York, D., Pedersen L. (1993) Particle mesh Ewald: An N·log(N) method for Ewald sums in large systems. J Chem Phys 98:10089–10092.

86. Hess B, H. Bekker, H.J.C. Berendsen, J.G.E.M. Fraaije (1997) INCS: A linear constraint solver for molecular simulations. J. Comput Chem 18:1463–1472.

