## Supplementary material for "Tension mediated mechanical activation and pocket delipidation lead to an analogous MscL state": Suppl_Tables and Figures_Wang_Lane et al_Pliotas_2021

This file contains supplementary:

Materials and methods
Figures with legends 1 to 10
Tables with legends 1 to 2
Legend for supplementary Video 1

A

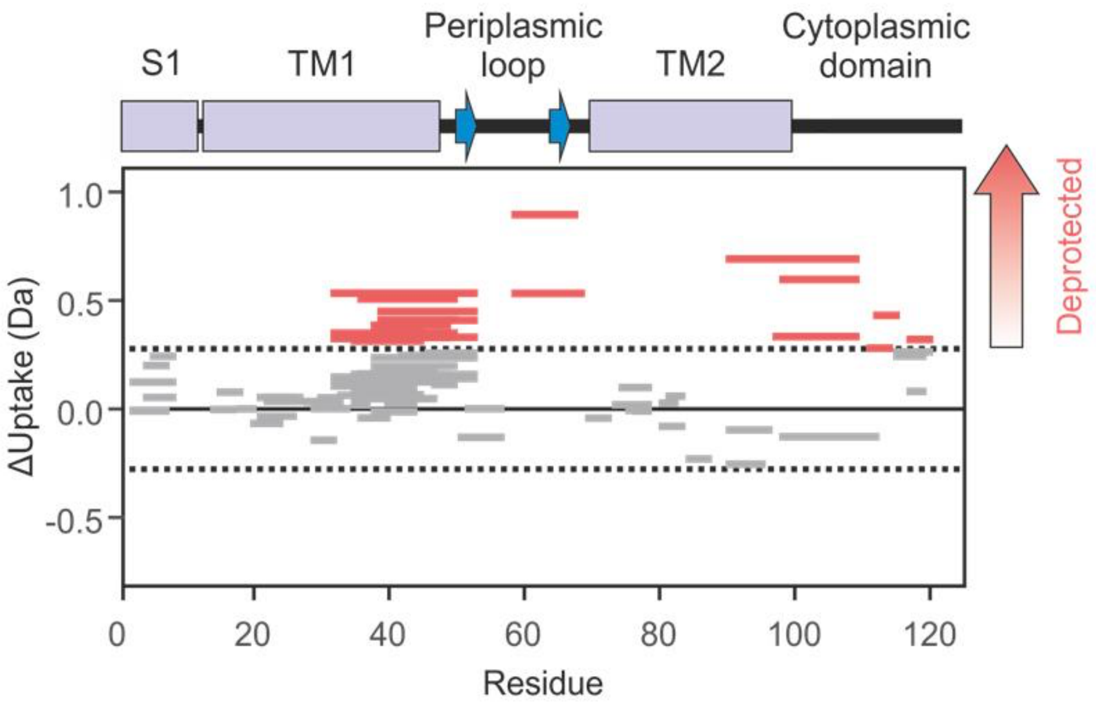


B

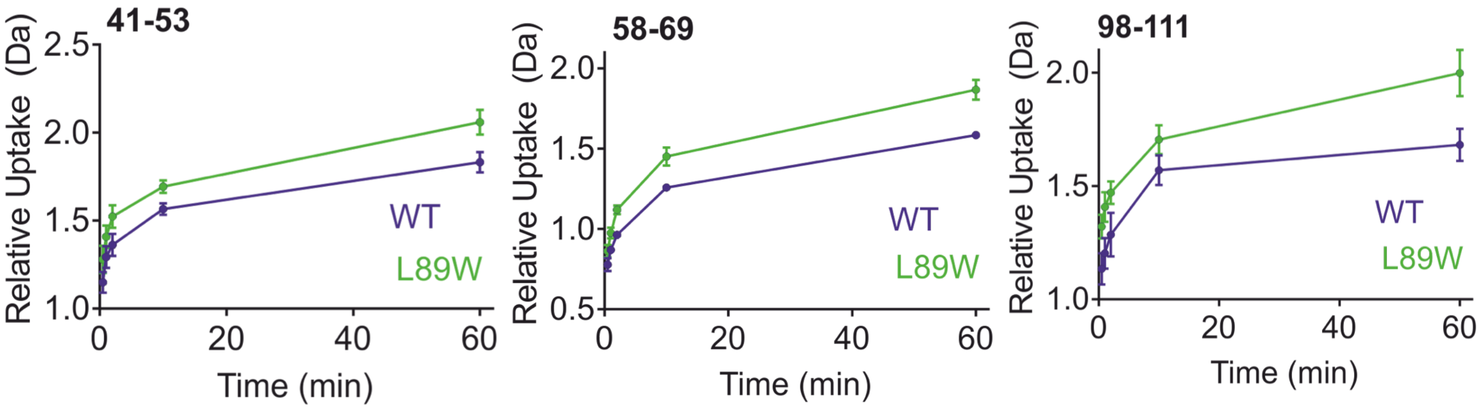


**Supplementary Figure 1.** A. Wood's plots showing the summed differences in deuterium uptake in MscL over all five HDX timepoints, comparing wildtype MscL with L89W MscL (Wood's plots were generated using Deuteros(76)). Peptides coloured in red, are deprotected from exchange in L89W MscL. No peptides were significantly protected from exchange in L89W MscL compared with wild type MscL. Peptides with no significant difference between conditions, determined using a 95% confidence interval (dotted line), are shown in grey. B. Example deuterium uptake curves for MscL WT (blue) L89W (green). The residue numbers in each peptide are indicated in the top left of each plot.

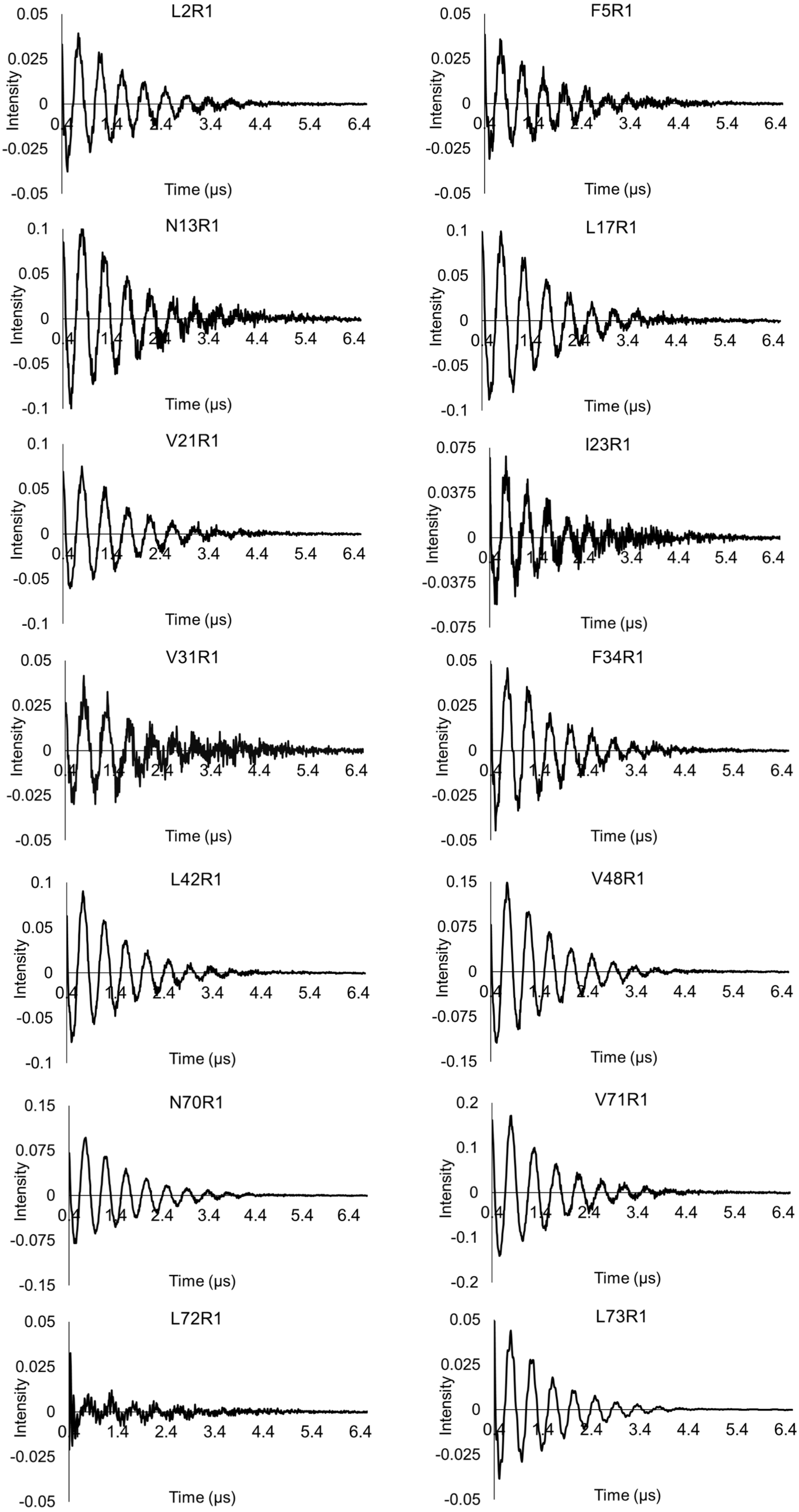
L2R1
F5R1
N13R1
L17R1
V21R1
I23R1
V31R1
F34R1
L42R1
V48R1
N70R1
V71R1
L72R1
L73R1
Intensity
Time (µs)
0.4
1.4
2.4
3.4
4.4
5.4
6.4
0.05
0.025
0
-0.025
-0.05
0.1
-0.1
0.075
0.0375
-0.0375
-0.075
0.15
-0.15
0.2
-0.2

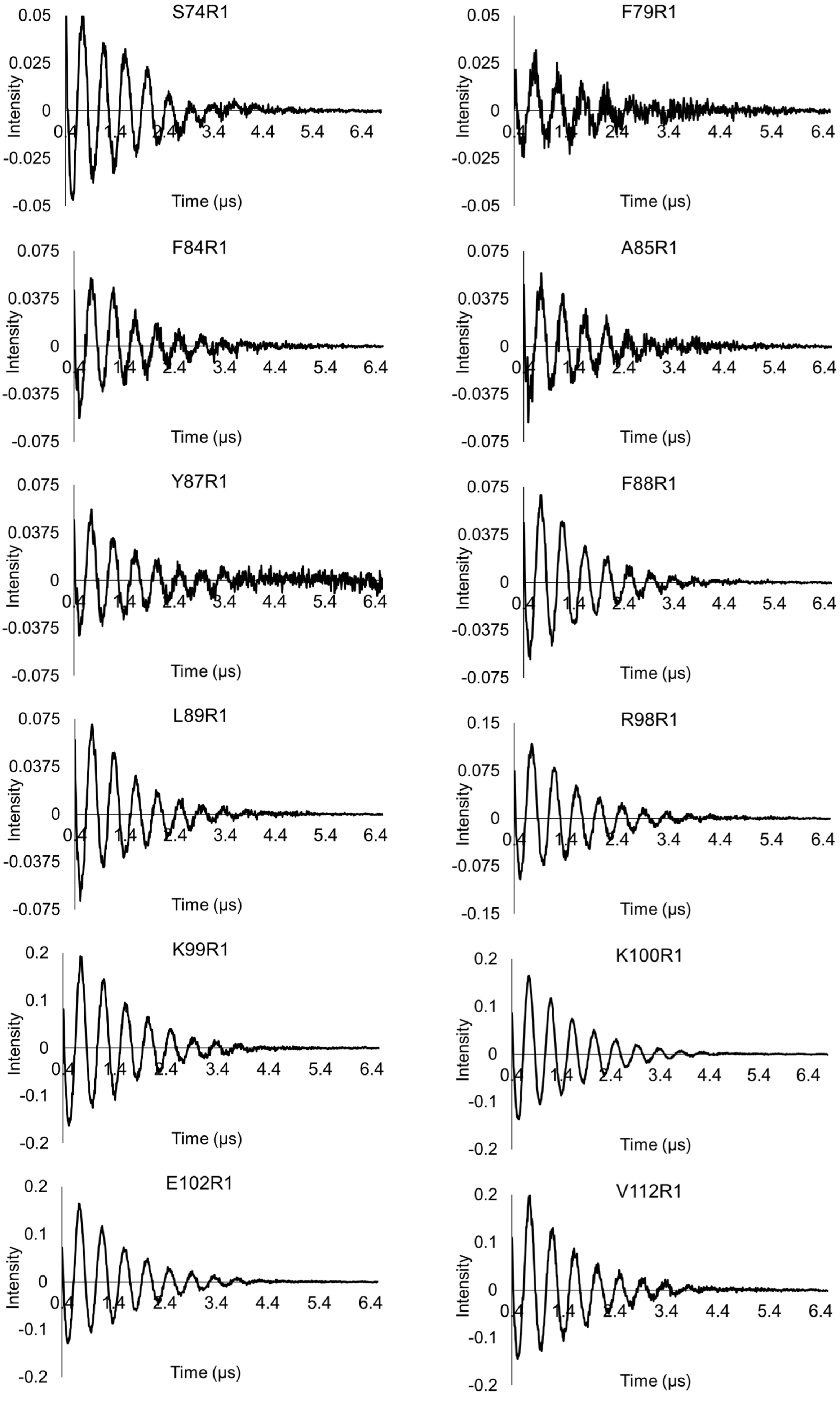


**Supplementary Figure 2.** Background corrected time-domain 3pESEEM raw experimental spectra used for the solvent accessibility determination of spin labelled TbMscL residues.

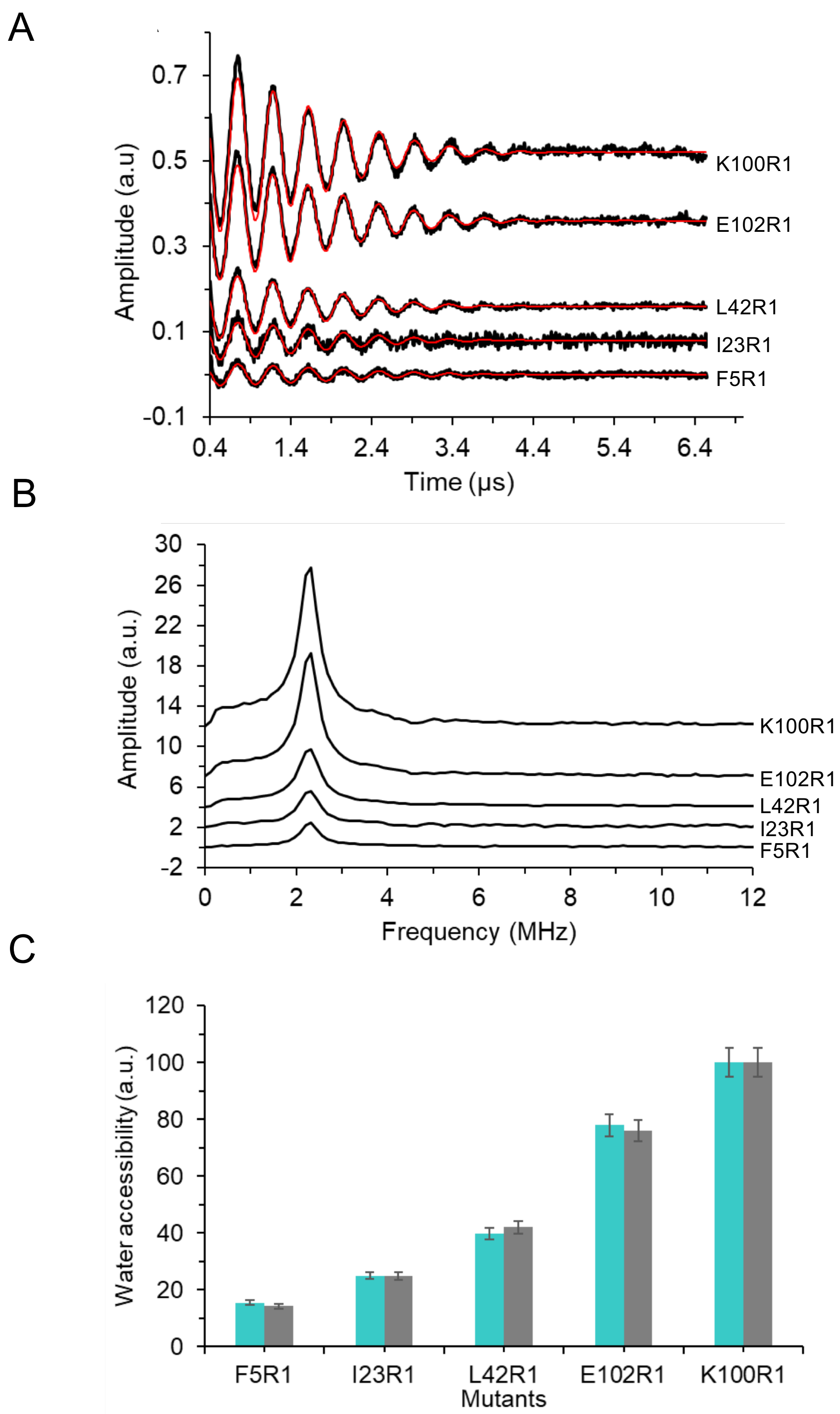


**Supplementary Figure 3.** A Background-corrected time-domain 3pESEEM raw spectra (black traces) with fitting (red) of representative in respect of solvent exposure level and MscL domain coverage spin-labelled mutants. F5R1 is found on the S1, I23R1 and L42R1 on TM1, and K100R1 and E102R1 are at the interface between TM2 and the CHB. B Frequency domain spectra of 3pESEEM data of F5R1, I23R1, L42R1, K100R1, and E102R1. C Column bar charts representing solvent accessibility parameters obtained by two different analysis method approaches. For each sample, the cyan bars correspond to the solvent accessibility derived from the deuterium amplitudes in frequency domain 3pESEEM spectra and normalized to the highest accessibility corresponding to 100%. The grey bars correspond to the solvent accessibility determined from the fitting model to the time domain 3pESEEM spectra and normalized to the highest accessibility corresponding to 100%. More details on data processing in *Materials and Methods.*

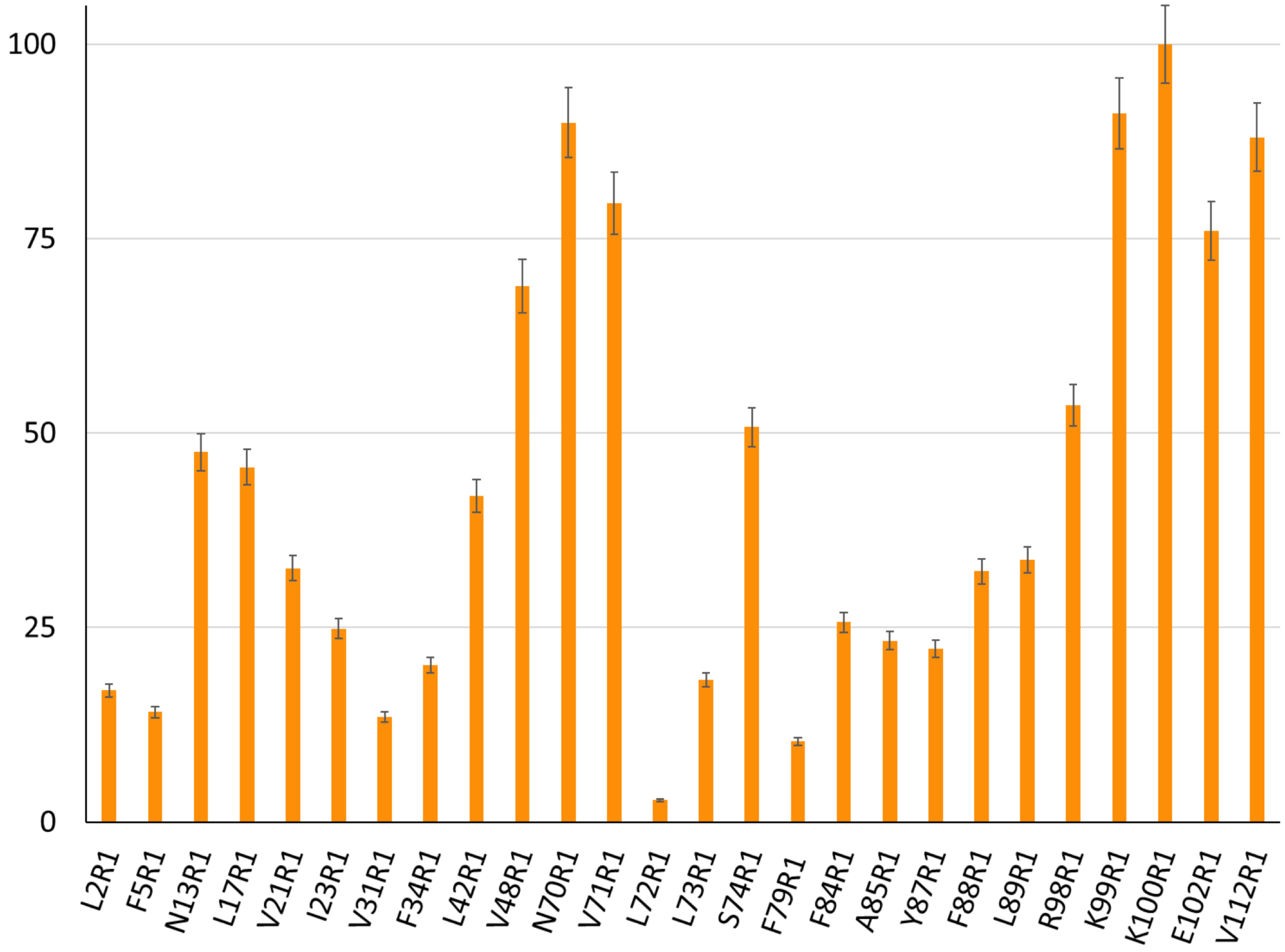


**Supplementary Figure 4.** The column bar charts represent the deuterium (or solvent) accessibility of each spin labelled residue derived from the fitting of 3pESEEM time-domain traces, normalised on a scale between 0 and 100**.** Errors are calculated at 5% to compensate for fitting errors and differences in the relaxation times of different spin labelled residues.

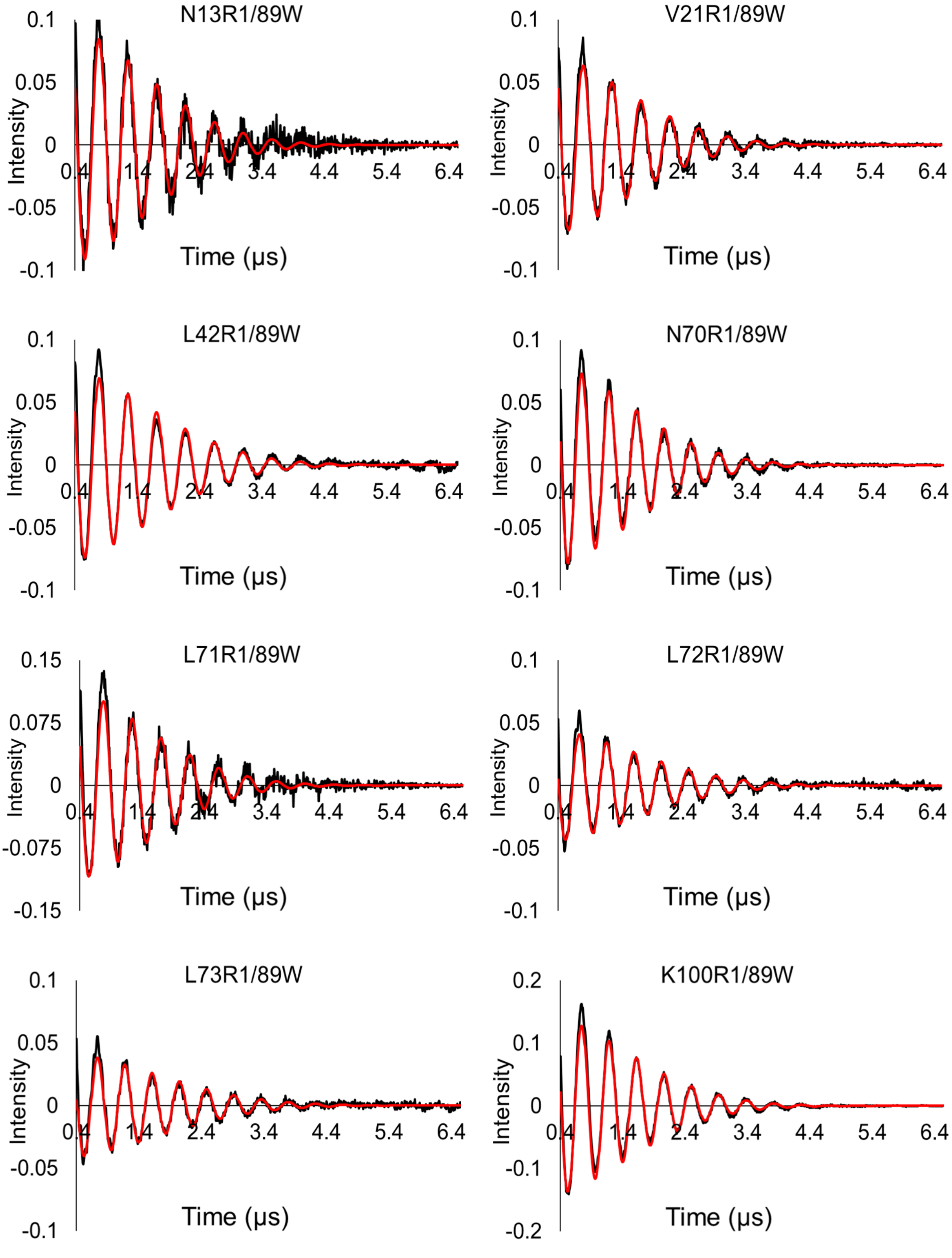


**Supplementary Figure 5.** Time-domain 3pESEEM raw (black) and fitted (red) experimental spectra of the double TbMscL mutants N13R1/89W, V21R1/89W, L42R1/89W, N70R1/89W, L71R1/89W, L72R1/89W, L73R1/89W, and K100R1/89W.

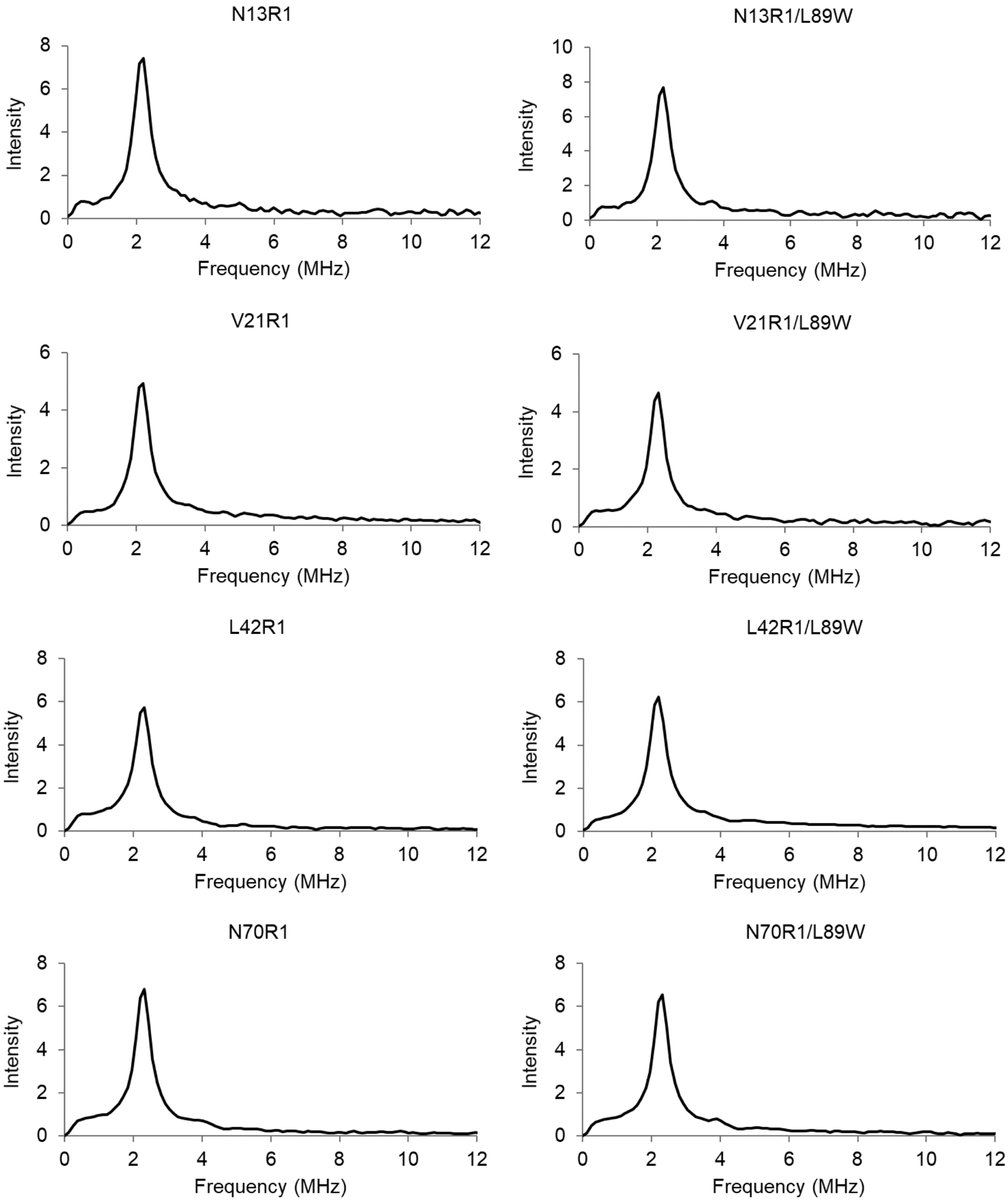

N13R1
N13R1/L89W
V21R1
V21R1/L89W
L42R1
L42R1/L89W
N70R1
N70R1/L89W
Intensity
Frequency (MHz)
0
2
4
6
8
10
12

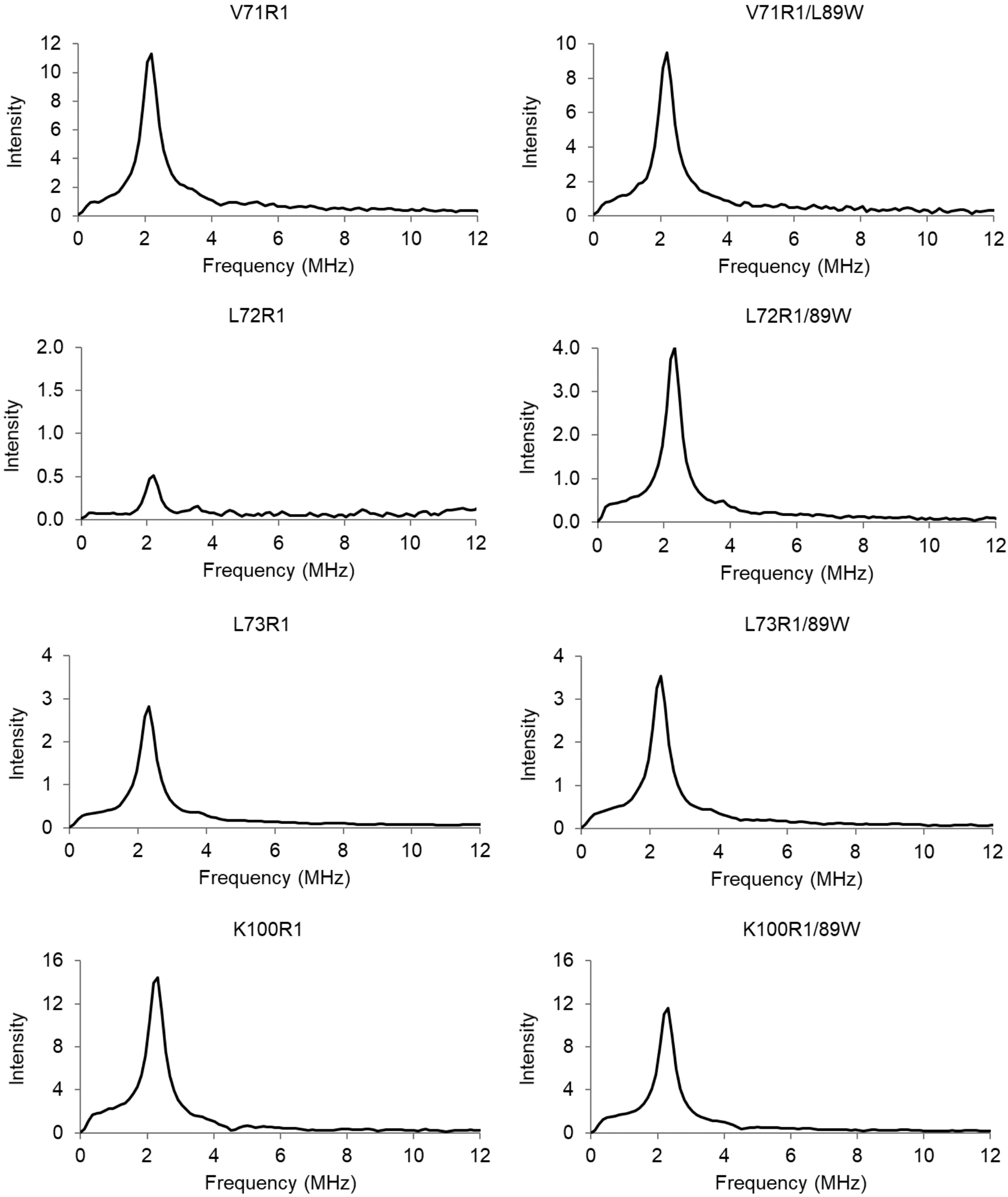


**Supplementary Figure 6.** Frequency domain spectra of 3pESEEM data used for solvent accessibility determination of TbMscL spin labelled residues 13R1, 13R1/89W, 21R1, 21R1/89W, 42R1, 42R1/89W, 70R1, 70R1/89W, 71R1, 71R1/89W, 72R1, 72R1/89W, 73R1, 73R1/89W, 100R1, and 100R1/89W.

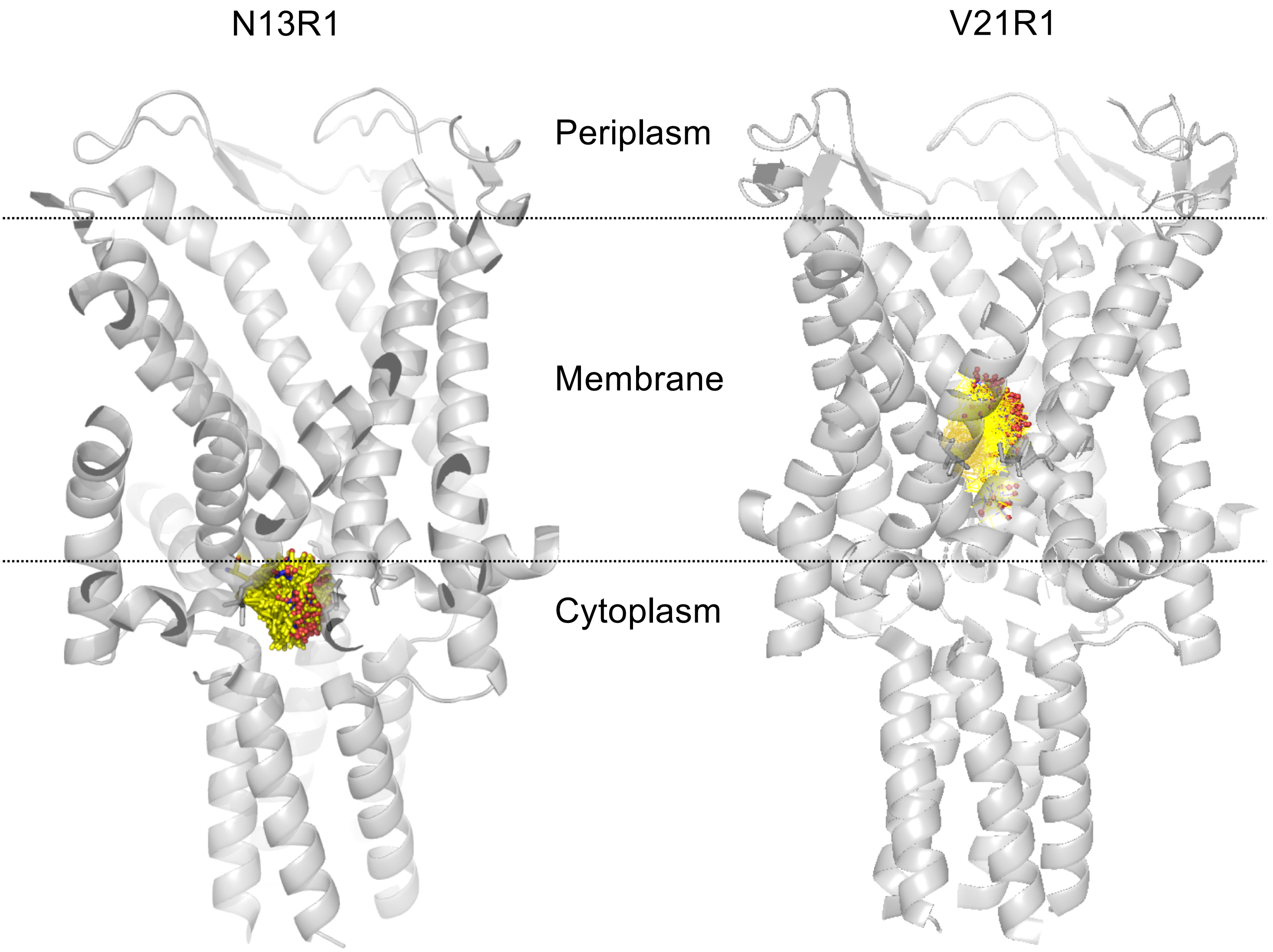


**Supplementary Figure 7.** *In sillico* spin labelling of TbMscL (PDB 2OAR, closed state) N13 and V21. N13R1's side chain is already solvent exposed in the closed state, due to pointing into the cytoplasm, thus unaffected in the expanded intermediate state (Fig 3).

A

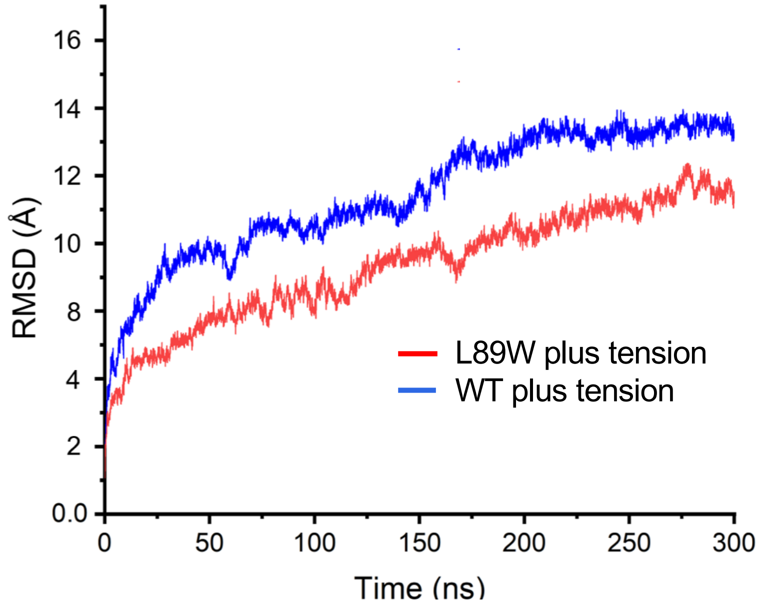


B

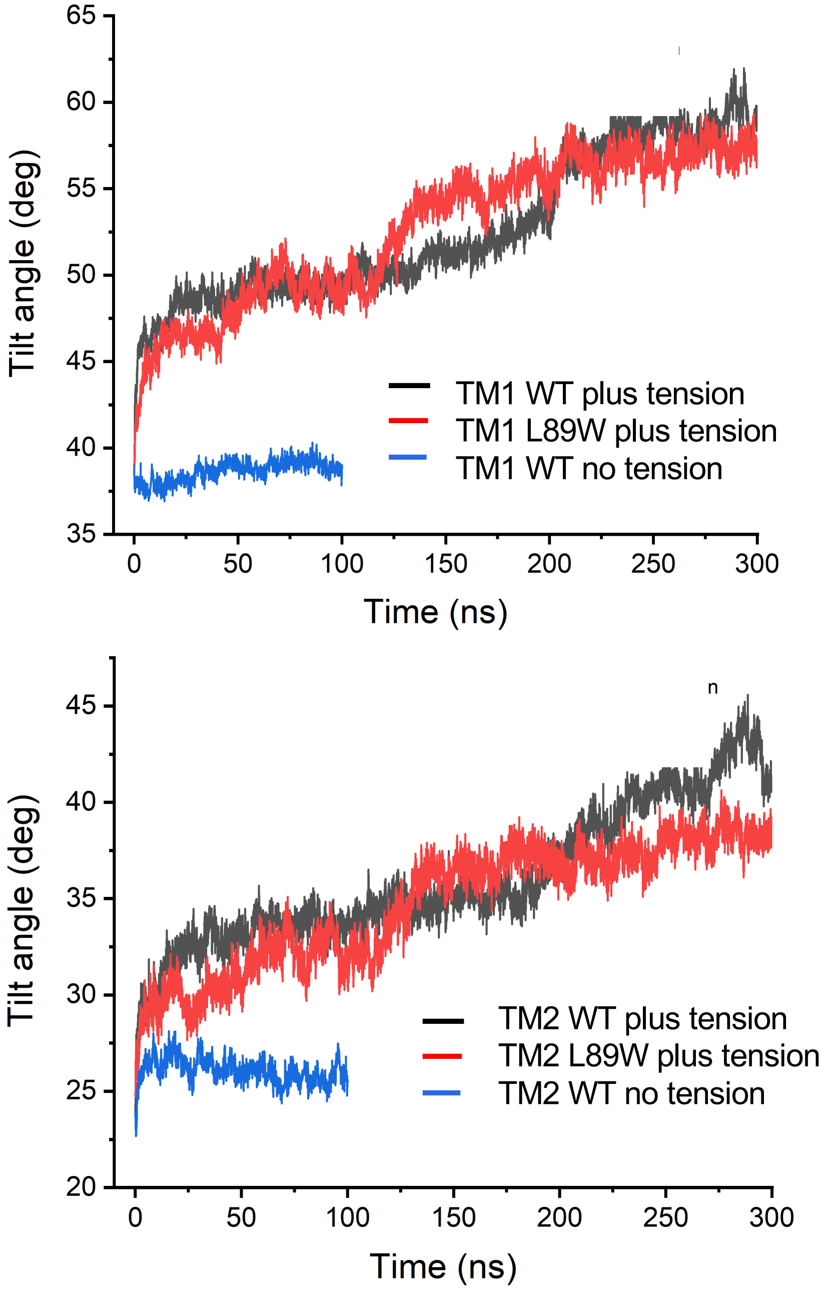


**Supplementary Figure 8.** A. RMSD between WT and L89W TbMscL under applied to the x-y, parallel to the membrane plane bilayer tension. B. Tilting angle comparison between under tension and no tension and modified states**.** Tilting angle of TM1 (left) and TM2 (right) with respect to the Z-axis over time.

A

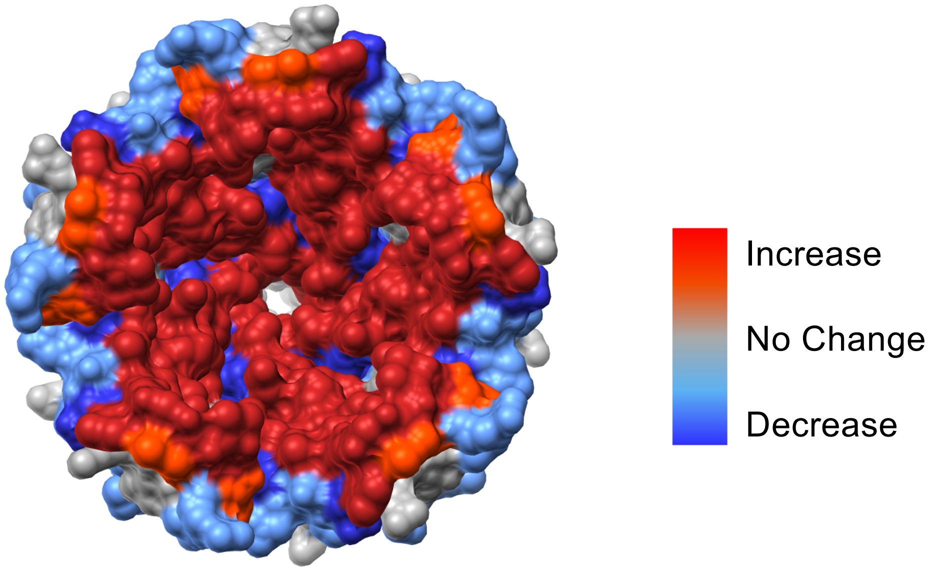


B

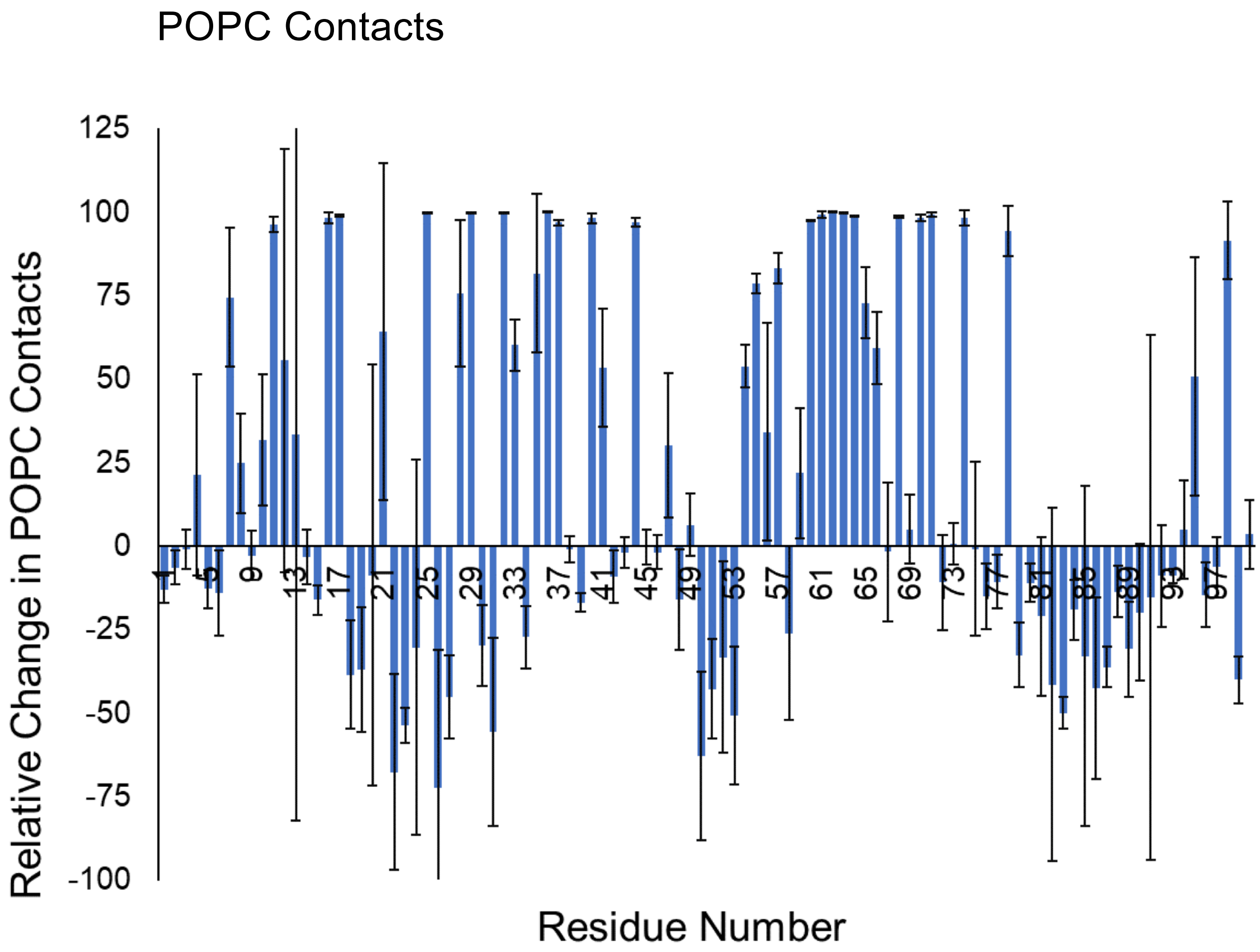


**Supplementary Figure 9.** A. MscL top pore view showing relative changes in the number of lipid contacts following simulated tension application in the membrane during MD. The blue regions show decrease in lipid contacts, while the red regions show an increase in lipid contacts. B. relative difference between the expanded tension-activated state and the closed (no tension) state of POPC contacts with TbMscL residues.

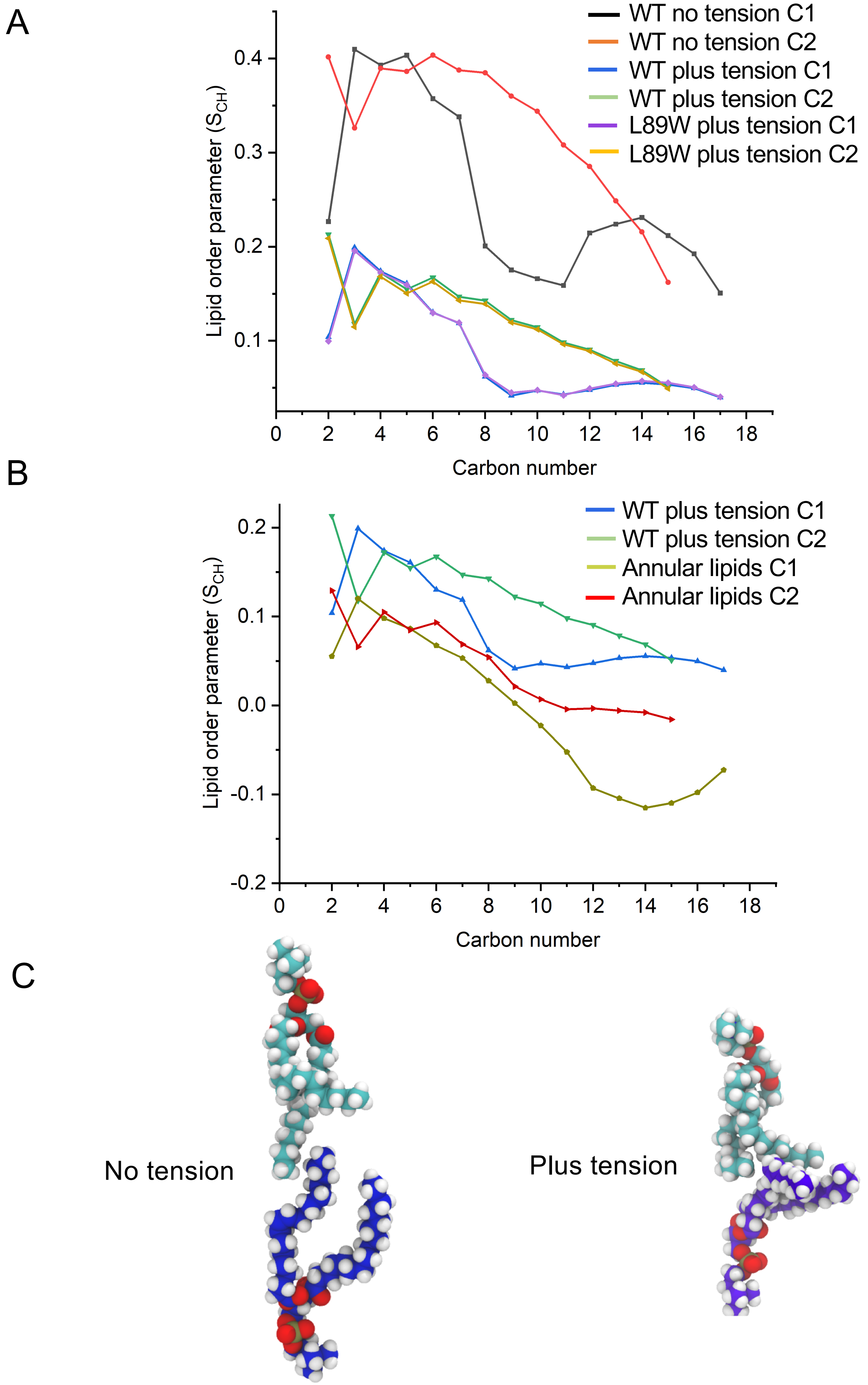


**Supplementary Figure 10.** Comparison of the lipid order parameters A. for all the lipids included in the simulation of WT MscL with no tension, WT and L89W MscL under tension. Lipid chains are more horizontally oriented under tension compared to chains with no tension while in both L89W and WT MD simulations lipid chains adopt similar horizontal orientations. B. for all bilayer and annular lipids included in the WT MscL MD simulations under tension. Annular lipids are more "horizontally" oriented than bulk bilayer lipids. C. representative lipids in MD with and no tension. $S_{ch}$ was calculated using the formula: $S_{ch} = ½ * (3 \cos^2θ – 1)$, where *θ* is the angle formed between the (C-1) and (C+1) vector and the membrane horizontal axis.

**Supplementary Table 1.** HDX Data Summary Table
SD = standard deviation, CI = confidence interval

| **Data Set** | **Tb MscL** | **L89W MscL** |
|---|---|---|
| **HDX reaction details** | 50 mM potassium phosphate pH 7.4, 300 mM NaCl, 0.05% DDM | |
| **HDX time course (min)** | 0.5, 1, 2, 10, 60 | |
| **HDX control samples** | Maximally-labelled controls were not performed. | |
| **Back-exchange** | ~ 30 % | |
| **# of Peptides** | 101 | 101 |
| **Sequence coverage** | 83 % | 83 % |
| **Average peptide length / Redundancy** | 7.3 / 5.6 | 7.3 / 5.6 |
| **Replicates (biological or technical)** | 3 technical, 2 biological | 3 technical, 2 biological |
| **Repeatability** | 0.049 (average SD) | 0.053 (average SD) |
| **Significant differences in HDX (delta HDX > X D)** | Reference | 99% or 95 % CI in summed data using Deuteros |

**Supplementary Table 2.** Overall RMSD among structures generated in this study by MD (under membrane tension) and the expanded x-ray MscL structure (PDB 4Y7J)

| | WT under tension (MD) (Å) | L89W under tension (MD) (Å ) | Expanded x-ray structure (Å) |
|---|---|---|---|
| WT under tension (MD) (Å) | - | 3.1 | 3.3 |
| L89W under tension (MD) (Å) | 3.1 | - | 3.4 |

**Supplementary Video 1. Gating transitions MscL under applied membrane tension**. WT TbMscL (PDB 2OAR) was used as the initial x-ray model (closed state) and tension was applied to the lipid bilayer over 300 ns. Side and top views in cartoon mode of MscL (light blue) and a single subunit (pale yellow). Lipids are not shown for clarity. Video was implemented using Chimera and frames from 1, 40, 80, 160 and 300 ns time points of the MD simulation.
